# Central nervous system–derived inflammasome cytokines drive neuroglial injury and cerebral oedema in viral encephalitis, despite corticosteroid therapy

**DOI:** 10.1101/2025.10.10.681150

**Authors:** Franklyn Nkongho Egbe, Cordelia Dunai, Claire Hetherington, Sarah A. Boardman, Laura Bricio Moreno, Bethany Facer, Cory Hooper, David Haw, Alexandra-Chloé Villani, Luca Lenzi, Sam Haldenby, Sally O. Oswald, Steve Patterson, Evelyn Kurt-Jones, Andrew David Luster, Claire E. Eyers, Stuart Allan, Mark A. Ellul, Dexenceph study group, Tom Solomon, Benedict D. Michael

## Abstract

Herpes simplex virus (HSV) is the leading cause of encephalitis, with high rates of morbidity and mortality. Dysregulated immune responses have been associated with poor outcomes in small series, driving interest in immunotherapies. However, the key mediators of neuropathology, their cellular sources, and responsiveness to corticosteroid treatment remain unknown.

We present findings from an RCT of dexamethasone in 55 adults with HSV encephalitis, alongside a representative murine model.

Clinical severity and adverse outcomes were associated with neuroglial injury biomarkers (GFAP, tau, UCH-L1), increased cerebral oedema on MRI, and higher IL-1RA, IL-18, and IL-6. These mediators were more abundant in matched CSF than serum, and were not reduced by dexamethasone. In the model, neurons, astrocytes and microglia robustly expressed IL-1 and IL-6, supporting their CNS origin.

These findings identify IL-1 and IL-6 signalling as key drivers of immunopathology in HSV encephalitis and potential therapeutic targets to mitigate neuroinflammation and improve outcomes.

**Funding:** National Institute for Health and Care Research (grant number 12/205/28).

## Introduction

Herpes simplex virus encephalitis is a severe infection and inflammation of the central nervous system (CNS) caused by herpes simplex viruses (HSV) and characterised by acute brain inflammation. Although the antiviral drug aciclovir improves outcomes from around 70% mortality, 10-20% of patients still die, and 30 – 70% develop long-term neurological sequelae, including cognitive deficits, behavioural difficulties, and seizures ^1,2^, imposing a substantial burden on patients, caregivers and healthcare systems^3^. The exact mechanism of HSV-induced neuroinflammation and brain injury remains unclear, yet elucidating these pathways is critical for identifying avenues for adjunctive therapy alongside antivirals to improve clinical outcomes.

Animal models of HSV encephalitis suggest that inflammatory cytokines and chemokines produced within the CNS play a vital role in both the control of viral replication and pathogenesis ^4–6^; the balance between viral control and host immune responses appears to influence clinical severity and outcome ^7,8^. Identifying the cellular origin and the pattern of inflammatory mediators driving neuroinflammation and brain injury may therefore provide insight into disease mechanisms and reveal targets for adjunctive immunotherapy which may require adequate CNS penetration.

To dampen the immune response, adjunctive treatment with dexamethasone, a broad immunomodulatory corticosteroid, is sometimes administered to patients with HSV encephalitis in clinical practice. However, a recent RCT of adjunctive dexamethasone showed that this did not improve clinical outcome overall ^9^. It is therefore critical to better understand the inflammatory processes and to determine which mediators are and are not modulated by corticosteroids. Therefore, identifying the array of mediators involved and determining how dexamethasone modulates them is essential not only for evaluating the drug’s efficacy but also for informing the development of alternative targeted immunotherapies.

Elevated levels of IL-6 and members of the IL-1 cytokine family have been associated with disease severity and poor outcomes in small series of HSV encephalitis patients ^10–13^, but their connection with HSV-induced neuroinflammation and brain injury is poorly defined. Furthermore, the role of IL-18, a member of the IL-1 superfamily, has not been characterised in human HSV encephalitis, despite experimental evidence that high levels of IL-18 and IL-1β, released via the NLRP3 inflammasome pathway, are associated with poor outcomes in murine models ^10,14–16^.

In this study of adults with HSV encephalitis, treated with aciclovir with or without intravenous dexamethasone, we characterised and defined the cellular origin of key inflammatory mediators associated with HSV-induced neuroinflammation and brain injury on neuroimaging, clinical severity and poor outcomes, examining the role of corticosteroids. We show that CNS-derived IL-1 family and IL-6 cytokines are associated not just with clinical severity but also with both serum and CSF neuroglial biomarker and radiological evidence of neuroinflammation and brain injury. Moreover, the levels of these key mediators were unaffected by dexamethasone treatment. Given the ineffectiveness of dexamethasone, we suggest that adjunct immunomodulatory therapy targeting the CNS-derived IL-6 and IL-1 family cytokines may provide an alternative effective treatment in HSV encephalitis adult patients.

## Results

### Interleukin-6 and interleukin-1 cytokines are associated with HSV encephalitis clinical severity and poor outcomes

CSF and blood samples were obtained from 55 participants (32 in the treatment group and 23 in the control group). The median ages of the treatment and control groups were 65 (interquartile range IQR: 56 - 73) years and 70 (IQR: 58 - 80) years, respectively. The groups had similar clinical characteristics and disease outcomes (Table 1).

**Table 1:** Characteristics of Participants.

| Characteristics | All participants<br>(n = 55) | Aciclovir +<br>Dexamethasone<br>Group<br>(n = 32) | Aciclovir only<br>Group<br>(n = 23) | P value |
| --- | --- | --- | --- | --- |
| Sex-Male | 25 (45.5) | 19 (59.3) | 11 (47.8) | 0.57 |
| Age* | 67 ( 57 - 76) | 65 ( 56 - 73) | 70 ( 58 - 80) | 0.19 |
| Clinical |  |  |  |  |
| CSF-WCC (per mm3)* | 76 ( 14.9 - 275.3 ) | 141 (33.5 - 288.0) | 36 (11.1 - 114.3) | 0.1 |
| CSF-RCC (per mm3)* | 29 (4.3 - 205.3) | 18 (5.0 - 116.0) | 32 (4.1 - 240.0) | 0.64 |
| CSF-protein (mg/dL)* | 18 (15.0 - 23.0) | 17.0 (15.0 - 18.0) | 22.5 (18.8 - 23.5) | 0.19 |
| Had seizures at admission | 25 (49.02) | 15 (50) | 10 (47.6) | 0.61 |
| HSV viral load (Log copies/mL)* | 4.66 (4.2 – 5.1) | 4.3 (4.0 – 5.1) | 4.7 (4.4 – 5.0) | 0.67 |
| Pre-treatment GCS |  |  |  |  |
| Normal (GCS = 15) | 27 (50) | 17(53.1) | 10(45.5) | 0.78 |
| Abnormal (GCS < 15) | 27 (50) | 15(46.9) | 12(54.5) |  |
| Outcome Scores |  |  |  |  |
| mRS (30 days) |  |  |  |  |
| Good Outcome (mRS<3) | 27 (51.9) | 17(56.7) | 8(36.4) | 0.24 |
| Poor Outcome (mRS>2) | 25 (48.1) | 13(43.3) | 14(63.7) |  |
| LOS (30 days) |  |  |  |  |
| Good recovery (LOS > 2) | 29 (58) | 18(64.29) | 11(50) | 0.47 |
| Poor recovery (LOS =2) | 21 (42) | 10(35.7) | 11(50) |  |
| GOS (26 weeks) |  |  |  |  |
| Good recovery (GOS =7\|8) | 15 (33.3) | 10(40) | 5(25) | 0.46 |
| Poor recovery (GOS <7) | 30 (66.7) | 15(60) | 15(75) |  |
| Barthel (30 days) |  |  |  |  |
| Independent (score =20) | 21(40.38) | 15(50) | 6(27.3) | 0.17 |
| High dependency (score <20) | 31(59.62) | 15(50) | 16(72.7) |  |

Summary statistics are **counts(column %)** or **\*median (IQR)**. **GCS**: Glasgow coma scale score, **mRS**: Modified Rankin Score, **LOS**: Liverpool Outcome Score, **GOS**: Glasgow Outcome Score, and **Barthel**: Barthel Index score

Four brain injury biomarkers and forty-eight cytokines, chemokines and growth factors were assessed in the serum and CSF of these patients using the Quanterix single-molecule array and Bio-Rad multiplex Luminex panels, respectively. The Glasgow coma scale (GCS) score at admission, where 15 is normal (less severe disease), and <15 indicates more severe disease, was used to assess clinical severity, while modified Rankin scale (mRS), Liverpool outcome score (LOS) and Barthel Index were used to assess the disability and functional outcomes either at discharge or 30 days post-admission.

Of the 48 cytokines, chemokines and growth factors assessed in CSF collected at admission, nine were associated with clinical severity, including the pro-inflammatory cytokines interleukin-1RA (*P* value = 0.003) and interleukin-18 (P value = 0.015) of the interleukin-1 superfamily, interleukin-12 (IL-12; p = 0.013), interferon-γ (IFNγ; p = 0.046), as well as leukocyte chemotactic chemokines, C-C motif chemokine ligand 3 (CCL3; p = 0.017), C-C motif chemokine ligand 5 (CCL5; *P* value = 0.003), C-C motif chemokine ligand 7 (CCL7; *P* value = 0.03), and granulocyte colony-stimulating factor (G-CSF; *P* value = 0.005). Most of the levels were significantly raised in patients with more severe disease (GCS score < 15), except for IL-12 and CCL5, where the levels were significantly higher in patients with a GCS score of 15. The ratios of IL-1α, IL-1β and IL-1RA were also associated with clinical severity (Fig. 1-a,c,d,f).

**Fig. 1.**
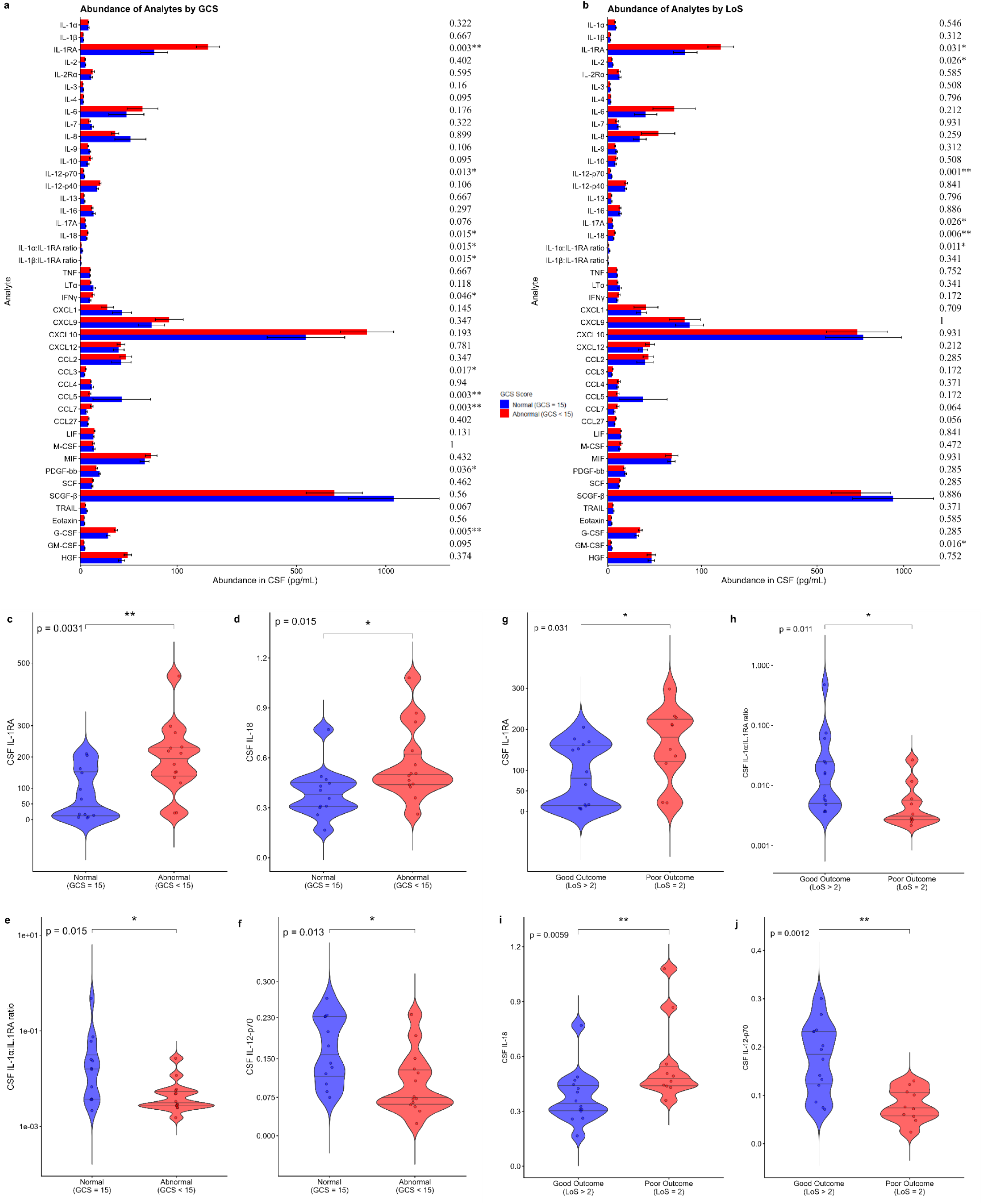
IL-1 inflammatory mediator levels in CSF are associated with clinical disease severity and poor outcomes following HSV encephalitis. **a,b,** Analyte abundance plots showing concentrations of mediators in CSF from encephalitis patients according to clinical severity and outcome, respectively. Each analyte represents median-centred and normalised values. *P* values are displayed on the right. **c–j,** levels of mediators in CSF stratified by GCS scores: **(c)** IL-1RA, **(d)** IL-18, **(e)** IL-1α:IL-1RA ratio, **(f)** IL-12; LOS: **(g)** IL-1RA ratio, **(h)** IL-1α:IL-1RA ratio, **(i)** IL-18, **(j)** IL-12. Means were compared using the Mann-Whitney test. The violin plots show the median and interquartile values; each data point represents one patient’s clinical sample. Disease severity is defined as a normal (GCS = 15) versus an abnormal (GCS < 15) Glasgow Coma Scale score. Clinical outcome is defined based on the Liverpool Outcome Score (LOS); a good outcome is LOS >2, and a poor outcome is an LOS of 2. * = *P* < 0.05, ** = *P* < 0.01, and *** = *P* < 0.001

Next, we determined which of these nine mediators were also associated with poor functional outcomes. We found that patients with poor outcomes (LOS = 2) had significantly higher CSF levels of IL-1RA (180.7 vs 81.0 pg/mL, p = 0.03) and IL-18 (0.48 vs 0.34 pg/mL; p = 0.01) compared to patients with better outcomes (Fig. 1-b,g,i and Supplementary Table 1); and both cytokines were significantly negatively correlated with LOS (IL-1RA: tau = -0.43, p =0.009; IL-18: tau = -0.42, p = 0.01) and mRS (IL-1RA: tau = 0.39, p = 0.01; IL-18: tau = 0.39, p = 0.01) (Extended Fig. 1-a,e,g). Although baseline CSF HSV viral load showed a positive correlation with IFN-γ, IL-6 and IL-10, it was not associated with clinical severity or outcome (Extended Fig. 1-a). Thus, only levels of IL-1RA and IL-18 in CSF appear to be associated with acute clinical severity and poor outcomes in adult patients with HSV encephalitis.

In serum, the pro-inflammatory cytokines, IL-1RA and IL-6, were associated with both clinical severity (significantly higher in patients with GCS scores <15) and poor outcomes (significantly higher in patients with LOS of 2, and mRS scores of 3 or more), (Fig. 2-a,c,e and Fig. 2-b,g,h). Both cytokines showed a negative correlation with GCS and LOS, and the Barthel Index score, reiterating their association with clinical severity and poor outcome (Extended Fig. 2). The levels of IL-1β:IL-1RA ratio, and antiviral cytokine TNF-related apoptosis ligand antagonist (TRAIL) were associated with milder disease (significantly higher in patients with a GCS score of 15), with TRAIL also significantly raised in patients with better outcomes (0.91 vs 0.17 pg/mL, p = 0.03), (Fig.2-a,b,d,f). Early aciclovir initiation was associated with improved outcomes (Supplementary Fig. 1)

**Fig. 2.**
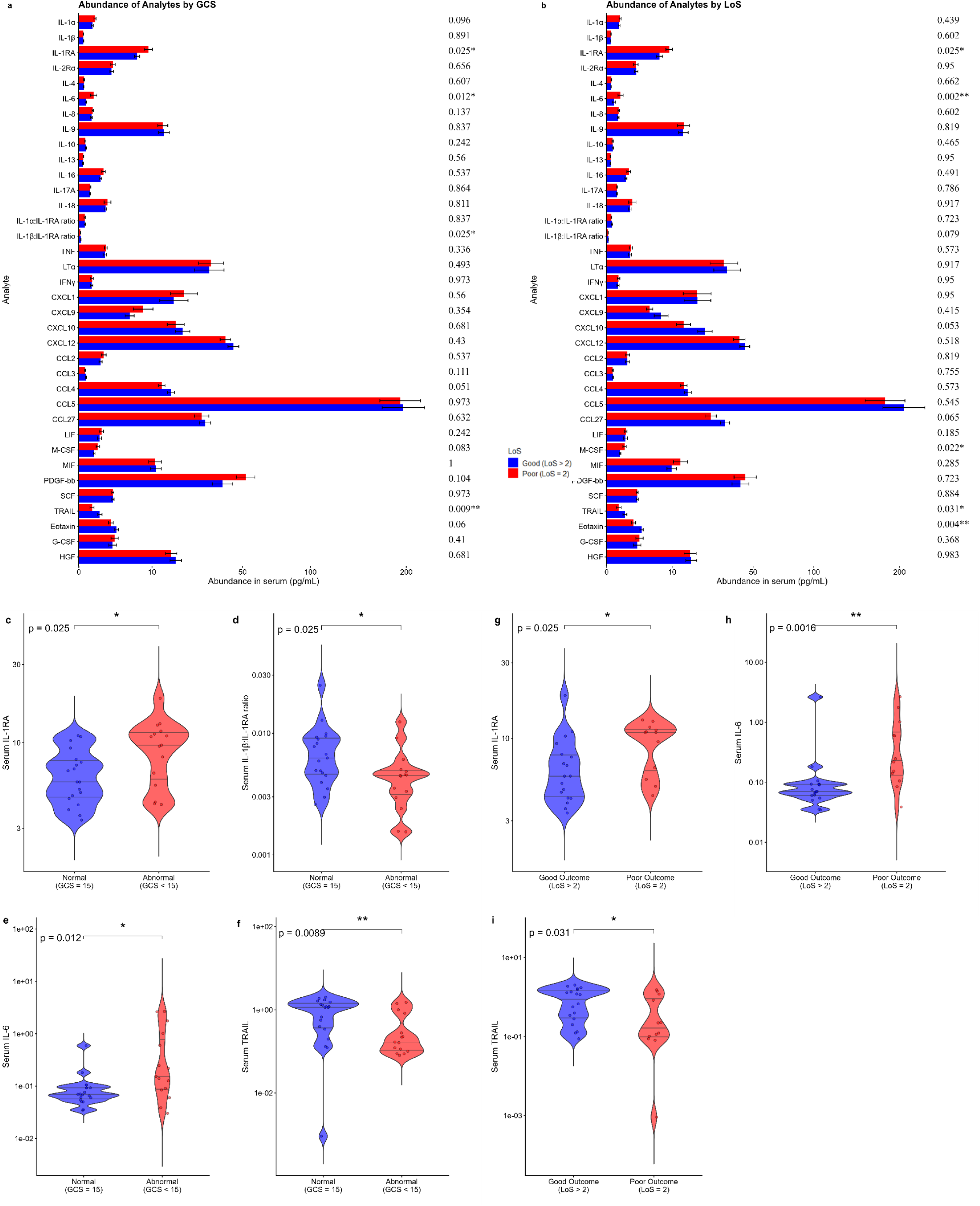
IL-6 and IL-1 inflammatory mediator levels in serum are associated with clinical severity and worse outcomes following HSV encephalitis. **a,b,** Analyte abundance plots showing concentrations of mediators in serum from encephalitis patients according to disease severity and outcomes, respectively. Each analyte represents median-centred and normalised values. *P* values are displayed on the right. **c–i**, Serum levels of mediators stratified by; **GCS**: **(c)** IL-1RA, **(d)** IL-1β:IL-1RA ratio **(e)** IL-6, **(f)** TRAIL. **LOS**: **(g)** IL-1RA **(h)** IL-6, **(i)** TRAIL. Means were compared using the Mann-Whitney test. The violin plots show the median and interquartile values; each data point represents one patient’s clinical sample. Disease severity is defined as a normal (GCS = 15) versus an abnormal (GCS < 15) Glasgow Coma Scale score. Clinical outcome is defined based on the Liverpool Outcome Score (LOS); a good outcome is LOS >2, and a poor outcome is an LOS of 2. * = *P* < 0.05, ** = *P* < 0.01, and *** = *P* < 0.001

Comparison of the concentrations of key mediators in 24 patients with matched acute CSF and serum samples generally showed higher levels in CSF than in serum samples, indicating that the mediators likely originated predominantly in the CSF (Extended Fig. 8).

Taken together, these results show that the levels of proinflammatory mediators, particularly IL-1 family cytokines in CSF and IL-6 in blood, were associated with both clinical severity and poor outcomes following HSV infection and were likely to originate from the CNS.

### IL-1 superfamily cytokines are associated with neuroglial injury and neuroinflammation in HSV encephalitis

Given the significantly increased levels of the predominantly proinflammatory cytokines, including cytokines of the interleukin-1 superfamily, in patients with severe clinical disease and poor outcomes, we hypothesised that these cytokines drive neuroglial injury and neuroinflammation in HSV encephalitis. We thus determined if these cytokines were also associated with levels of brain injury biomarkers and with the volume of cerebral oedema on MRI scans. Spearman correlation analysis revealed positive associations between CSF concentrations of IL-1RA, IL-18, and IL-10 with GFAP, a biomarker of astrocytic damage, and an inverse association with CSF levels of IL-7, IL-12 and TRAIL (Fig. 3-a,b,c,e). We also found significant associations between CSF IL-18 with Neurofilament Light chain (NfL) - a biomarker for neuroaxonal damage and neurodegeneration (Fig. 3-d). There were also significant associations between CSF C-X-C motif chemokine ligand 9 (CXCL9), CXCL12, and CCL7 with GFAP (Fig. 3-e).

**Fig. 3.**
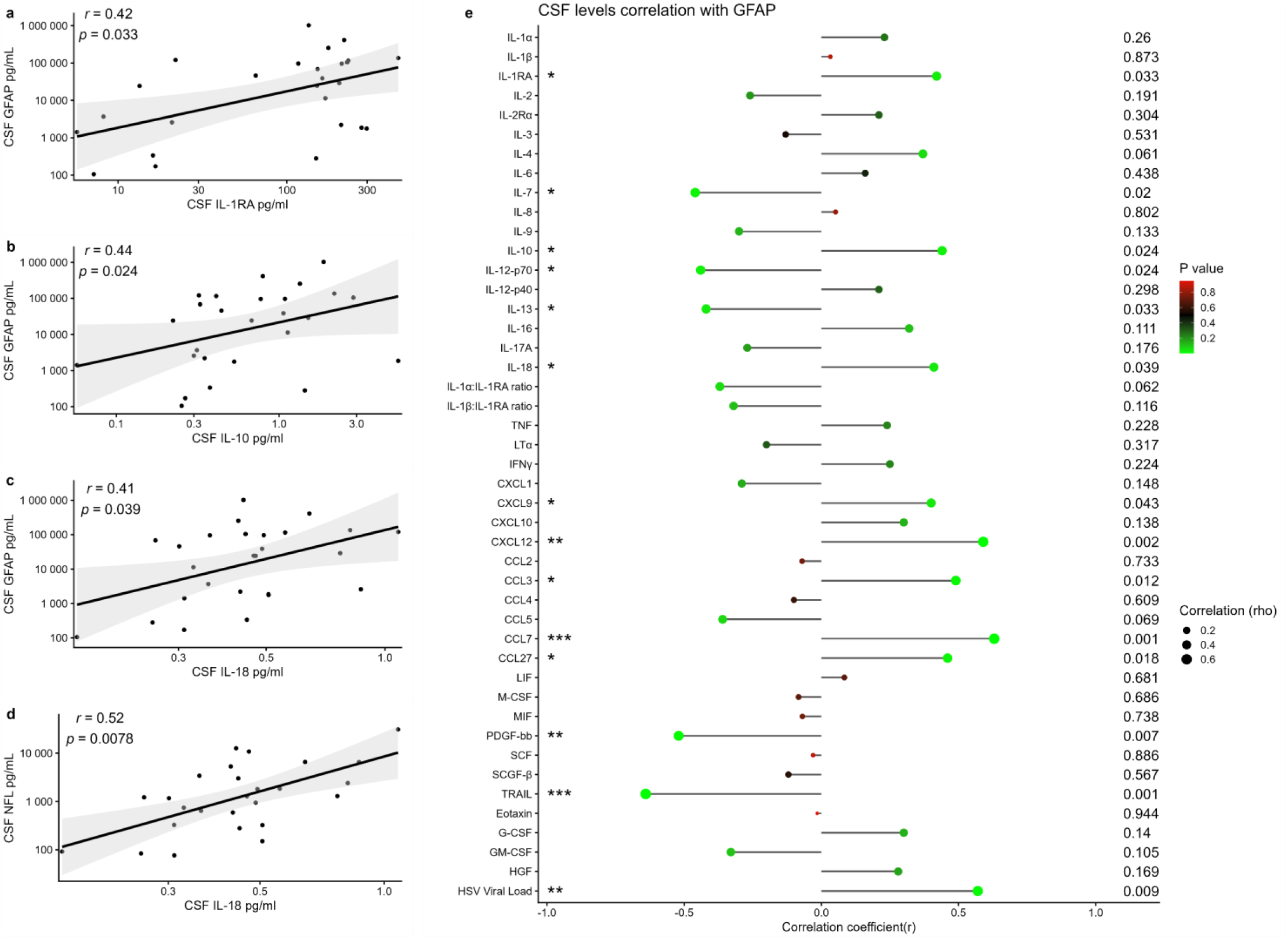
CSF levels of Cytokines of the Interleukin-1 superfamily are associated with neuroglial injury in HSV encephalitis. **a-d,** Spearman correlation between the concentration of mediators and biomarkers of neuroglial injury in CSF. **(a)** IL-1Ra, **(b)** IL-10, **(c)** IL-18 and **GFAP**, a marker of astrocytic damage; **(d)** IL-18 and **NfL**, a biomarker for neuroaxonal damage and neurodegeneration. The shaded areas represent the 95% confidence intervals (CIs). Each data point represents one patient. **e,** Forest plot of Spearman correlations between mediators and GFAP in **CSF**. Dot size corresponds to the Spearman correlation coefficient, while colour represents the P value. * = *P* < 0.05, ** = *P* < 0.01, and *** = *P* < 0.001

In serum, the levels of IL-1RA, IL-6, and M-CSF were positively associated with GFAP (Fig. 4) and IL-1RA and M-CSF with NfL (Extended Figure 5, Supplementary Table 1). A correlation analysis of all 38 participants with MRI scans and CSF samples, irrespective of when the CSF was collected, showed a positive association between CSF IL-18 and vasogenic oedema (Fig. 5-a,b), and CSF IL-1RA and cytotoxic oedema (Fig. 5-c,d); indicating the potential role of the inflammasome in the HSV-induced neuroinflammation. Similarly, we found a significant positive association between CSF IL-18 levels and cytotoxic oedema among 10 participants who had their admission MRI scans within three days of CSF collection (Fig. 5e and Extended Fig. 6). We did not find any positive association between these key mediators levels and cerebral oedema when we expanded the analysis to all 15 participants with MRI scans at admission (Extended Fig. 3), possibly because of the wide time range (1 – 9 days) between lumbar puncture and MRI scans.

**Fig. 4.**
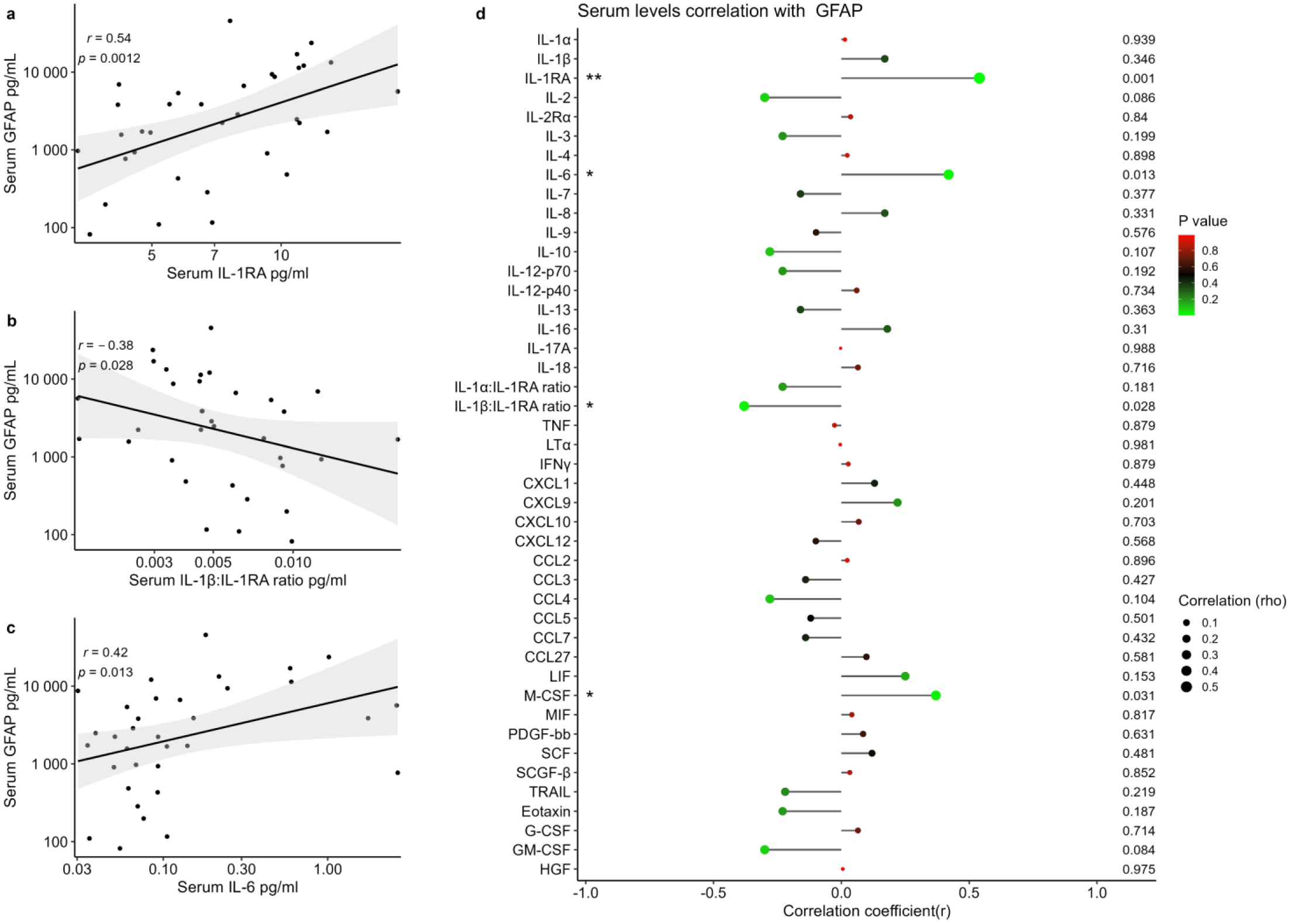
Blood levels of cytokines of the Interleukin-1 superfamily and IL-6 are associated with neuroglial injury in HSV encephalitis. **a-c,** Spearman correlation between the concentration of mediators and GFAP, a marker of astrocytic damage in Serum. **(a)**, IL-1Ra **(b)**, IL-1α:IL-1RA ratio **(c)**, IL-6. The shaded areas represent the 95% confidence intervals (CIs). **d**, Forest plot of Spearman correlations between mediators and GFAP in **Serum**. Each data point represents one patient. Dot size corresponds to the Spearman correlation coefficient, while colour represents the P value. * = *P* < 0.05, ** = *P* < 0.01, and *** = *P* < 0.001

**Fig. 5.**
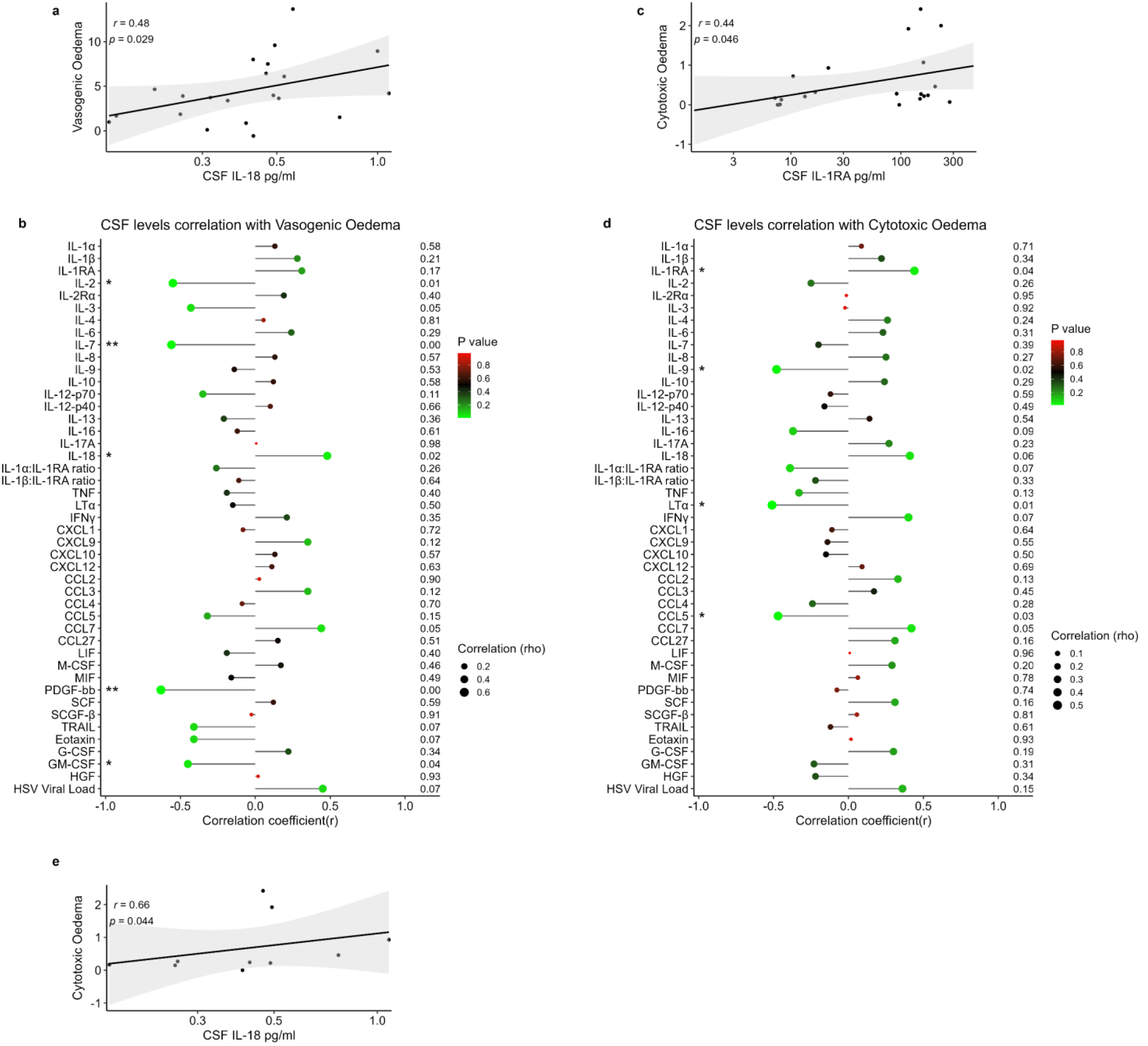
Cytokines of the Interleukin-1 superfamily are associated with neuroinflammation in HSV encephalitis. **a, c,** Spearman correlation between CSF concentration of mediators and MRI scans for all 38 participants with an *MRI scan and CSF sample collected at any time point*. **(a)**, IL-18 **(c)**, IL-1RA. The shaded areas represent the 95% confidence intervals (CIs). **b,d**, Forest plot of Spearman correlations between mediators and MRI scans. Each data point represents one patient. Dot size corresponds to the Spearman correlation coefficient, while colour represents the P value. **e**, Spearman correlation between CSF concentration of mediators and Cytotoxic oedema for **10 participants** with an MRI scan and CSF sample collected within 3 days. * = *P* < 0.05, ** = *P* < 0.01, and *** = *P* < 0.001

### Serum levels of IL-1RA, IL-6 and M-CSF associated with outcome in HSV encephalitis

We applied statistical modelling to identify key inflammatory mediators in serum and CSF that could predict good clinical outcomes in HSV encephalitis. For each sample type, a pre-specified subset of biologically plausible predictors was selected to avoid overfitting in this small dataset. For serum, IL-1RA, IL-6, M-CSF, and CCL2 were chosen based on exploratory analysis and prior literature. For CSF, the subset included IL-1RA, IL-1α, IL-1β, IL-18, and CCL3.

Outcomes were binarized using clinically relevant thresholds: mRS ≤ 1 (favourable), GCS of 15 (favourable), and length of stay ≤ 2 weeks (short). We used the ridge regression model, after evaluation (with and without an interaction term between dexamethasone use and time since randomisation) and cross-validation to account for the limited sample size and multicollinearity. Across all outcomes, serum-based models outperformed CSF-based models in terms of cross-validated AUC. Notably, serum IL-1RA, IL-6, and M-CSF emerged as significant predictors of outcome. These associations remained robust under bootstrapping, with narrow confidence intervals and low p-values. In contrast, none of the CSF cytokines showed statistically significant associations with outcome, and VIFs indicated substantial multicollinearity (e.g., VIF > 6 for CCL3), limiting interpretability.

### Changes in mediator levels in CSF and blood were unrelated to adjunctive dexamethasone treatment

To determine whether adjunctive dexamethasone treatment was associated with changes in analyte levels, we compared concentrations of key analytes in matched longitudinal samples obtained up to 9 days (early samples) and at day 10 or later post-randomisation (late samples) within each randomised group: aciclovir-only and aciclovir plus dexamethasone (Fig. 6). A ten-day cut-off was chosen based on the evidence from the primary analysis that participants who received dexamethasone before 10 days had better outcomes compared to those who received it later ^9^. Of the 55 participants, 14 had matched CSF and 20 had matched serum samples. No significant changes in analyte levels were observed upon adjunctive dexamethasone treatment (i.e. in the aciclovir plus dexamethasone group), suggesting that dexamethasone treatment did not influence the levels of these cytokines. However, we noted that CSF and serum levels of IL-1RA and IL-6 decreased significantly in the aciclovir only group (p < 0.05) (Fig. 6-a), while CSF IL-1α:IL-1RA, serum IL-1α:IL-1RA and IL-1β:IL-1RA ratios increased significantly (p <0.05), changes not observed in the aciclovir plus dexamethasone group (Fig. 6-a).

**Fig. 6.**
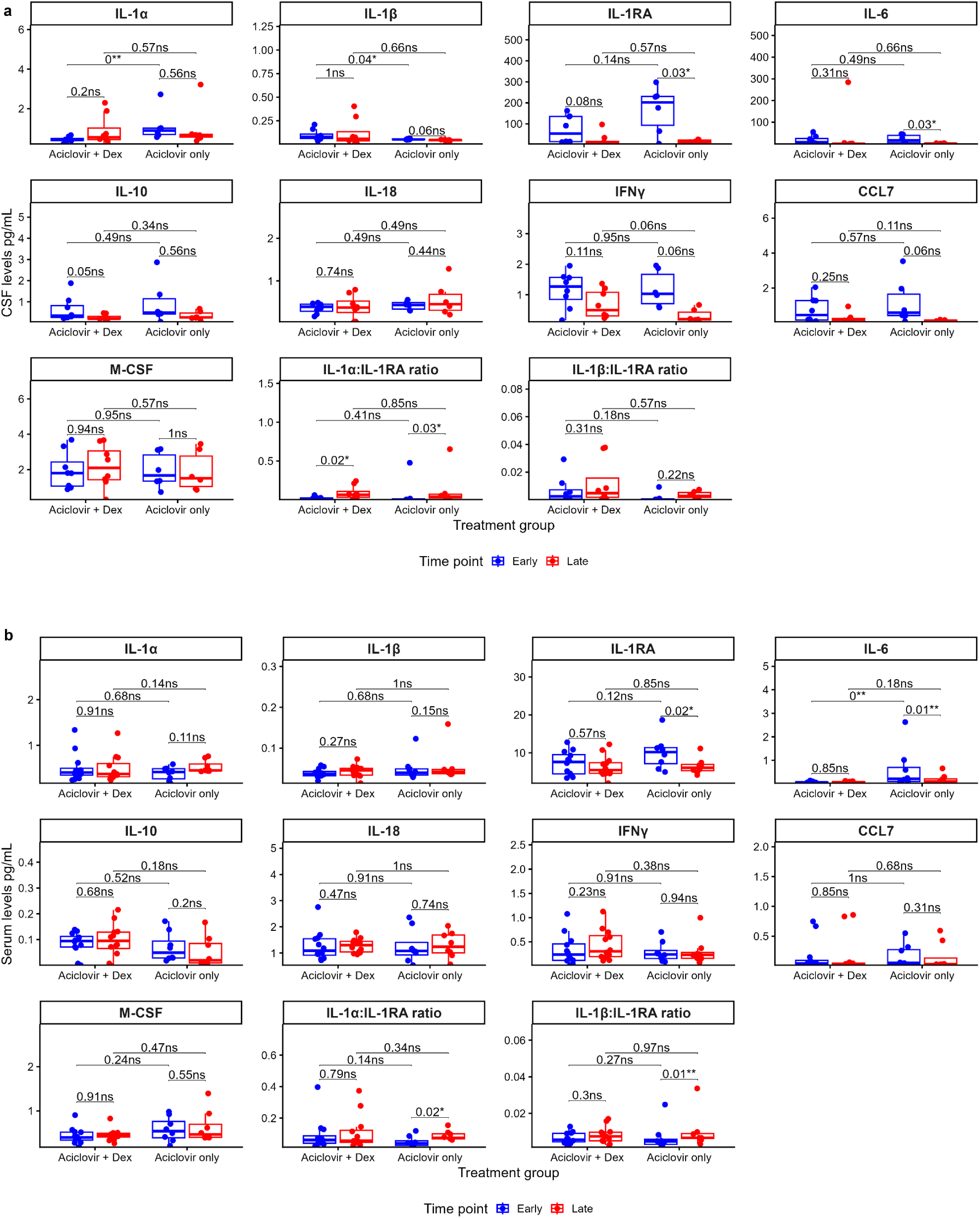
Changes in CSF and blood levels of mediators were unrelated to adjunct dexamethasone treatment. **a,b.** Boxplots comparing mediator levels in matched early and late clinical samples in both groups **(a) CSF:** 14 participants; 8 and 6 in the aciclovir + Dex and aciclovir only groups, respectively and (**b) Serum**: 20 participants; 12 and 8 in the aciclovir + Dex and aciclovir only groups, respectively. Matched samples within each group were compared using the Wilcoxon signed-rank test, and unpaired comparisons between groups using the Mann-Whitney U test. The boxplots show the median and interquartile values; each data point represents one patient’s clinical sample. Samples collected within 9 days of randomisation were defined as early, and samples collected on or after day 10 post-randomisation as late. For participants with multiple samples within a given time window, only the earliest sample was included in the analysis

We next compared analyte levels between the two randomised groups in the early (<10 days post-randomisation) and late (≥ 10 days post-randomisation) sample collection periods. In the late samples, concentrations did not differ significantly (p > 0.05) for any analyte, indicating that changes in the levels of analytes were unaffected by dexamethasone (Fig. 6 and Extended Fig 7-a,c). We noted that in the early samples, CSF IL-1α, serum IL-6, GFAP, NFL, and Tau were significantly lower in the aciclovir plus dexamethasone group (p < 0.01), while CSF IL-1β was higher (p < 0.05), but these differences were not observed in the late sample period (Fig. 6 and Extended Fig 7-c).

Furthermore, we compared fold-change patterns of these key analytes in both randomised groups and observed similar trajectories, with no adjunctive dexamethasone treatment-specific patterns (Extended Fig. 4 and Extended Fig. 7-b,d). Both groups exhibited significant reductions in serum GFAP and Tau from acute to post-acute stages (p < 0.01), whereas NFL slightly increased and remained sustained (p = 0.5) (Extended Fig. 7c).

Together, these results indicate that adjunctive dexamethasone treatment did not substantially influence changes in key mediators associated with disease severity and poor outcomes.

### KEGG pathway enrichment, gene ontology functional and STRING analyses reveal commonalities to inflammatory pathways and immune regulatory functions of key mediators

To gain biological insights into the potential pathways and functions of identified key mediators, Kyoto Encyclopaedia of Genes and Genomes (KEGG) pathway enrichment and STRING protein-protein interaction analyses were done.

Gene ontology enrichment analysis for KEGG Pathways of the nine mediators in CSF associated with clinical severity (GCS < 15; Table 2a) revealed commonalities with other non-viral inflammatory diseases including malaria and African trypanosomiasis, as well as viral-mediated Influenza A, but more generally with cytokine-cytokine receptor interaction and the JAK-STAT signalling pathway (Table 2b and Fig. 7-a). In agreement, functional enrichment analysis confirmed immune regulation as the most significant biological function across these 9 proteins (Supplementary Fig. 2), reiterating the vital role of proinflammatory mediators in HSV encephalitis.

**Fig. 7a.**
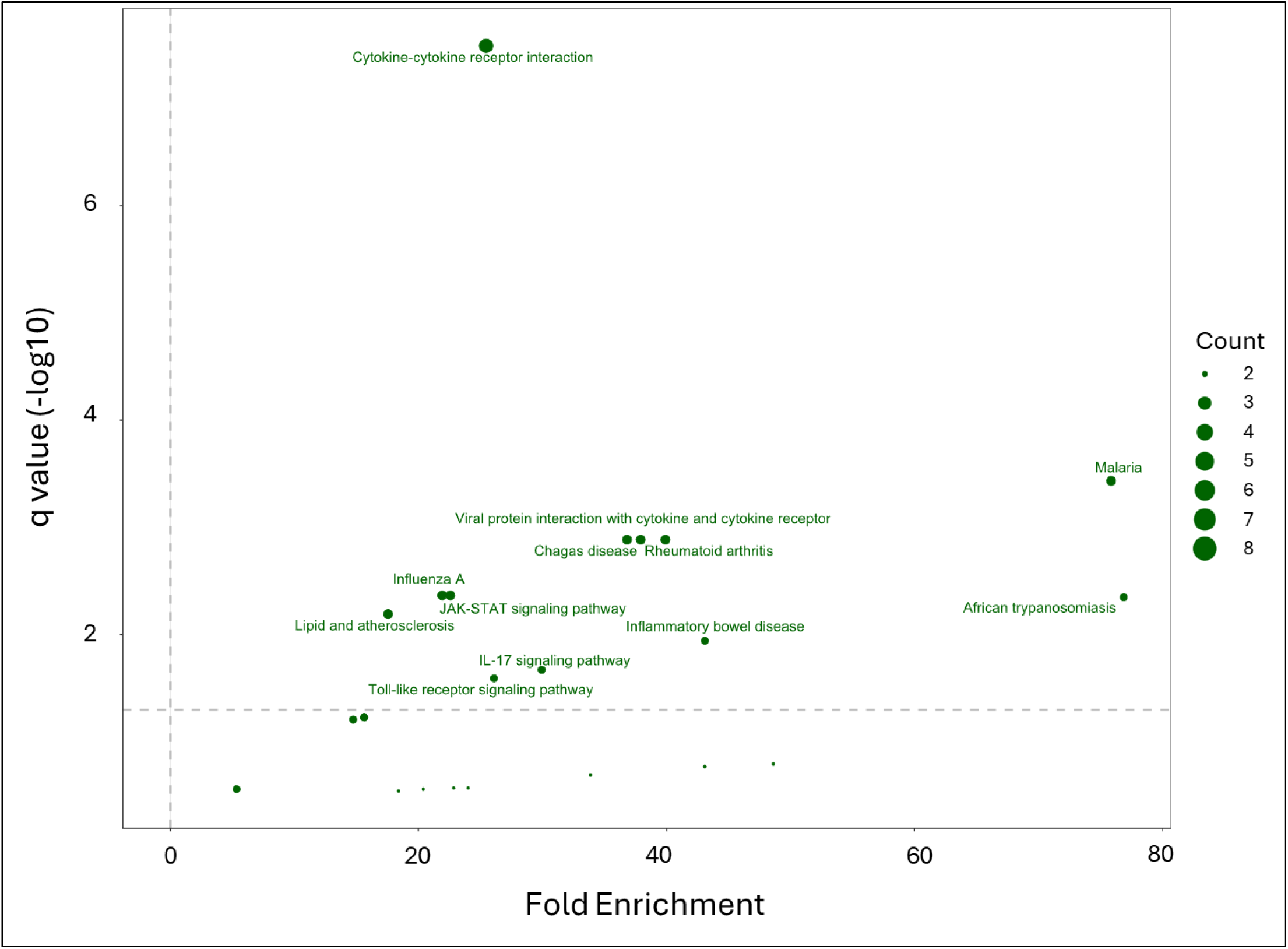
KEGG pathway enrichment analysis reveals commonalities to inflammatory pathways. Gene ontology enrichment analysis for KEGG Pathways of the 9 mediators associated with clinical severity was performed in DAVID functional annotation analysis. *P* values were adjusted for multiple hypothesis testing using the Benjamini-Hochberg correction and plotted against fold enrichment. The horizontal line represents an adjusted p-value of 0.05. Points are sized according to the number of proteins within each enriched term and coloured according to the enrichment category.

**Table 2.**
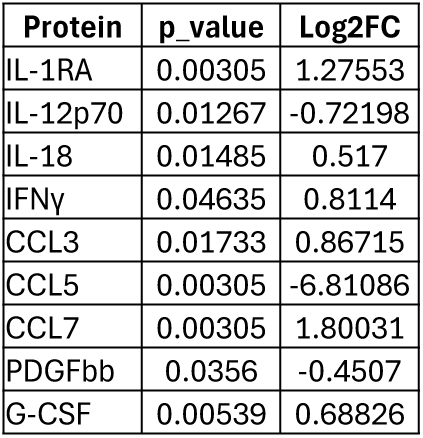
a: CSF mediators associated with clinical severity. **Clinical severity:** GCS<15/GCS=15

**Table 2b:**
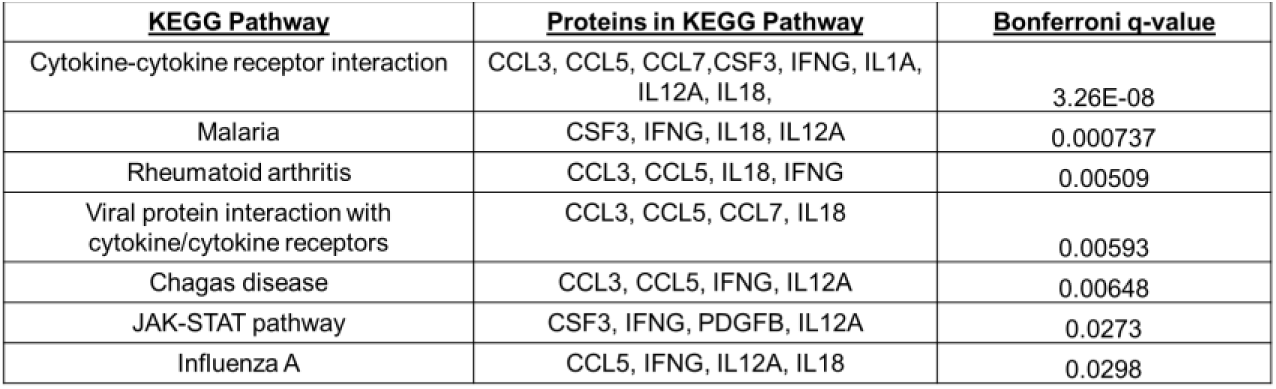
The top 7 associated KEGG pathways.

Network analysis of all 48 mediators assessed in CSF using STRING (v12) and KMEANS clustering to identify clusters of proteins with shared functions, produced three major clusters, viral protein interaction with cytokine/cytokine receptor, chemokine signalling, and cytokine-cytokine receptor interaction (Fig. 7-c), with key mediators identified in our study in the viral protein interaction with cytokine/cytokine receptor cluster.

**Figure 7-b,c.**
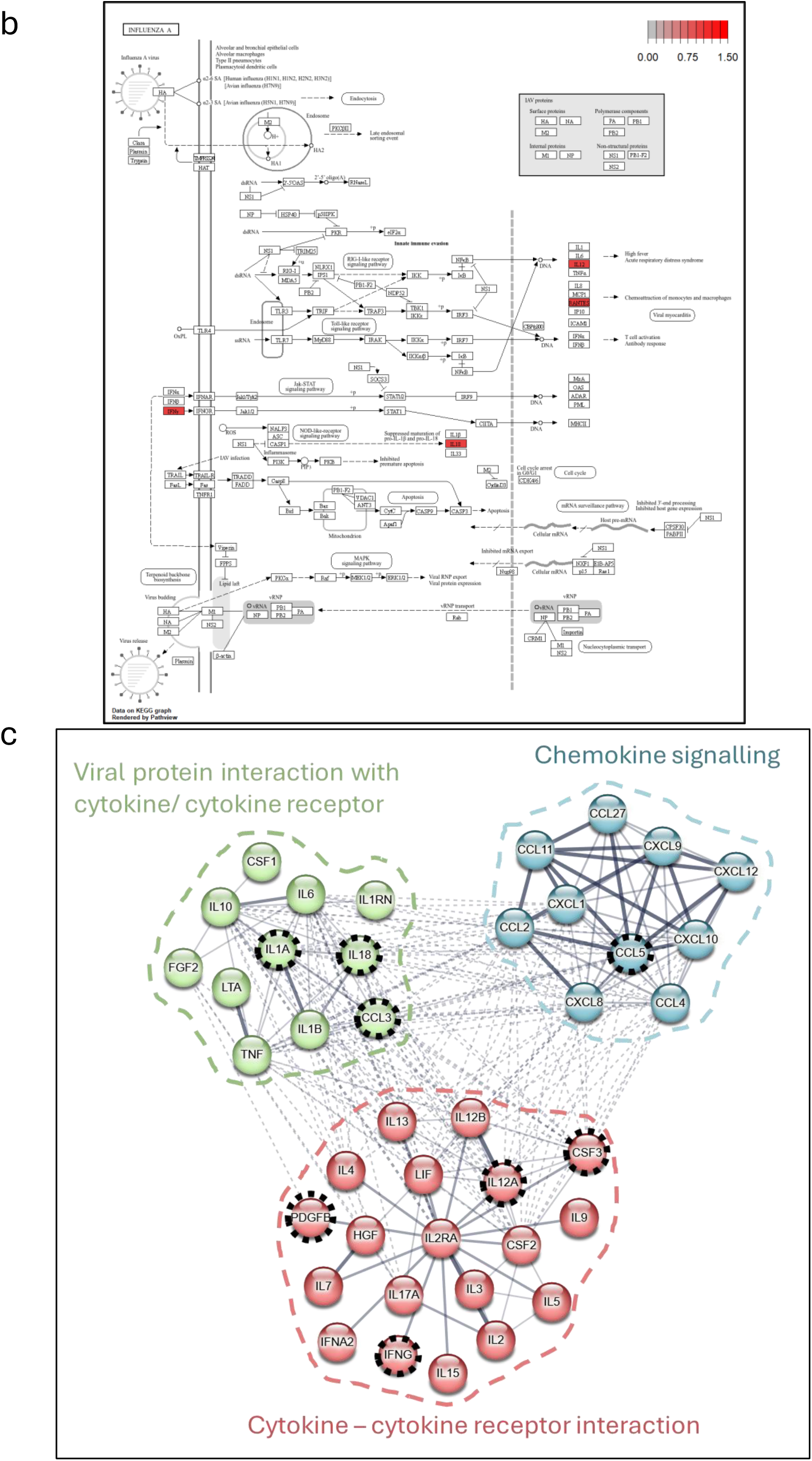
KEGG and STRING analyses highlight immune regulatory functions of key mediators. **b,** KEGG Influenza A pathway with highlighted study mediators. Cytokines identified in this study are mapped with colours indicating log₂ fold-change values, providing a visual representation of their regulation within biological contexts. The diagram was generated and visualised in R using the ‘pathview’ package (Luo & Brouwer, 2020). **c,** An interaction network of all measured mediators in CSF. Nodes arranged and colour-coded according to KEGG Pathways. The original network was generated in STRING, incorporating interactors supported by experimental or database evidence at medium confidence (interaction score ≥ 0.400). Line thickness reflects the strength of supporting data. Protein interactions that reached statistical significance (p < 0.05) are highlighted with a black dotted circular outline.

### HSV infection induces neuronal, microglial and astrocytic expression of IL-1β IL-6, and IL-18, and drove downstream inflammatory pathways in a murine model

Given the consistent associations between levels of IL-1 superfamily and IL-6 cytokines with clinical severity, poor outcomes, and neuroglial injury, and the greater abundance in CSF than serum, we hypothesised that expression of these cytokines would be upregulated specifically in neuronal and glial (astrocytes and microglial) following HSV infection. To test our hypothesis, we performed single-nuceli (sn) RNAseq to quantify heterogeneity of mRNA expression in different brain cell types subject to HSV infection, using an established HSV encephalitis murine model (Fig. 8a).^8^ Having confirmed the expression of HSV gene transcripts in various murine neuroglia, including neurons, microglia, and astrocytes (Fig. 8b), we next determined the transcript levels of the six cytokines most consistently identified in our clinical data: IL-1α, IL-1β, IL-1RA, IL-18, IL-6, and CCL3. We noted that in both the HSV-infected and control mice, these genes were most highly expressed in neurons, microglia, and astrocytes (Fig. 8c). Comparing their expression levels in cells from the HSV-1-infected and control mice, we noted in infected microglia, IL-18 exhibited significantly higher expression (log2fold change = 3.80, *P* value = 0.0056); in astrocytes, IL-1 and IL-6 (IL-1: log2fold change =0.94, *P* value < 0.001; IL-6: log2fold change =2.11, *P* value = 0.00012), and in neurons, IL-6 (log2fold change = 2.11, *P* value = 0.00012), (Fig. 8d). These data indicates that HSV infection induces neuronal, astrocytic, and microglial expression of IL-1, IL-6, and IL-18.

**Fig 8.**
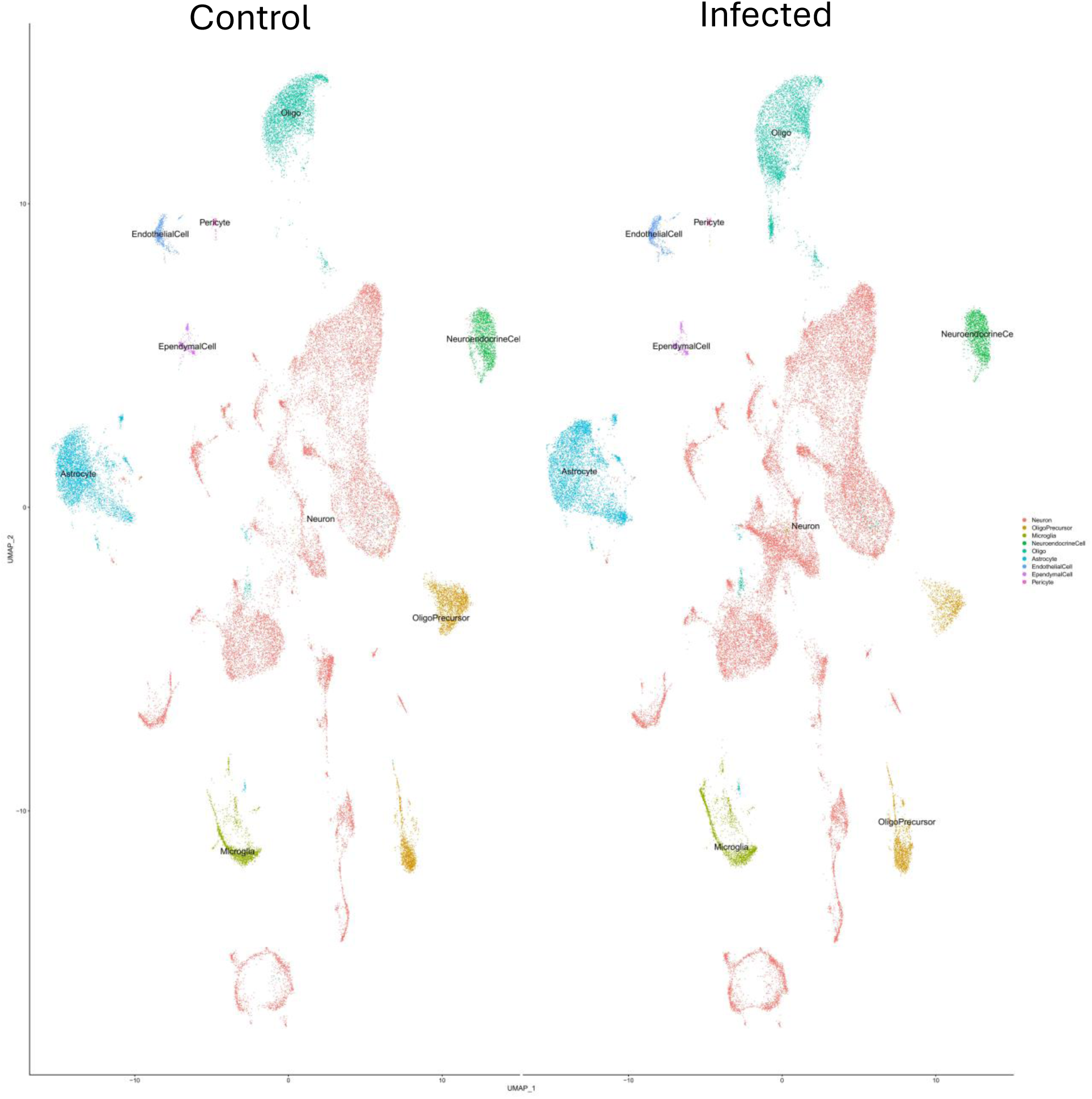
HSV infection induced neuronal and glial expression of IL-1, IL-6 and IL-18 in a murine model. **Fig 8a. UMAP plot of scRNAseq data**. Colours are mapped to the clusters (annotated on the right); broader cell-type definitions are overlayed on the plot

**Fig 8b.**
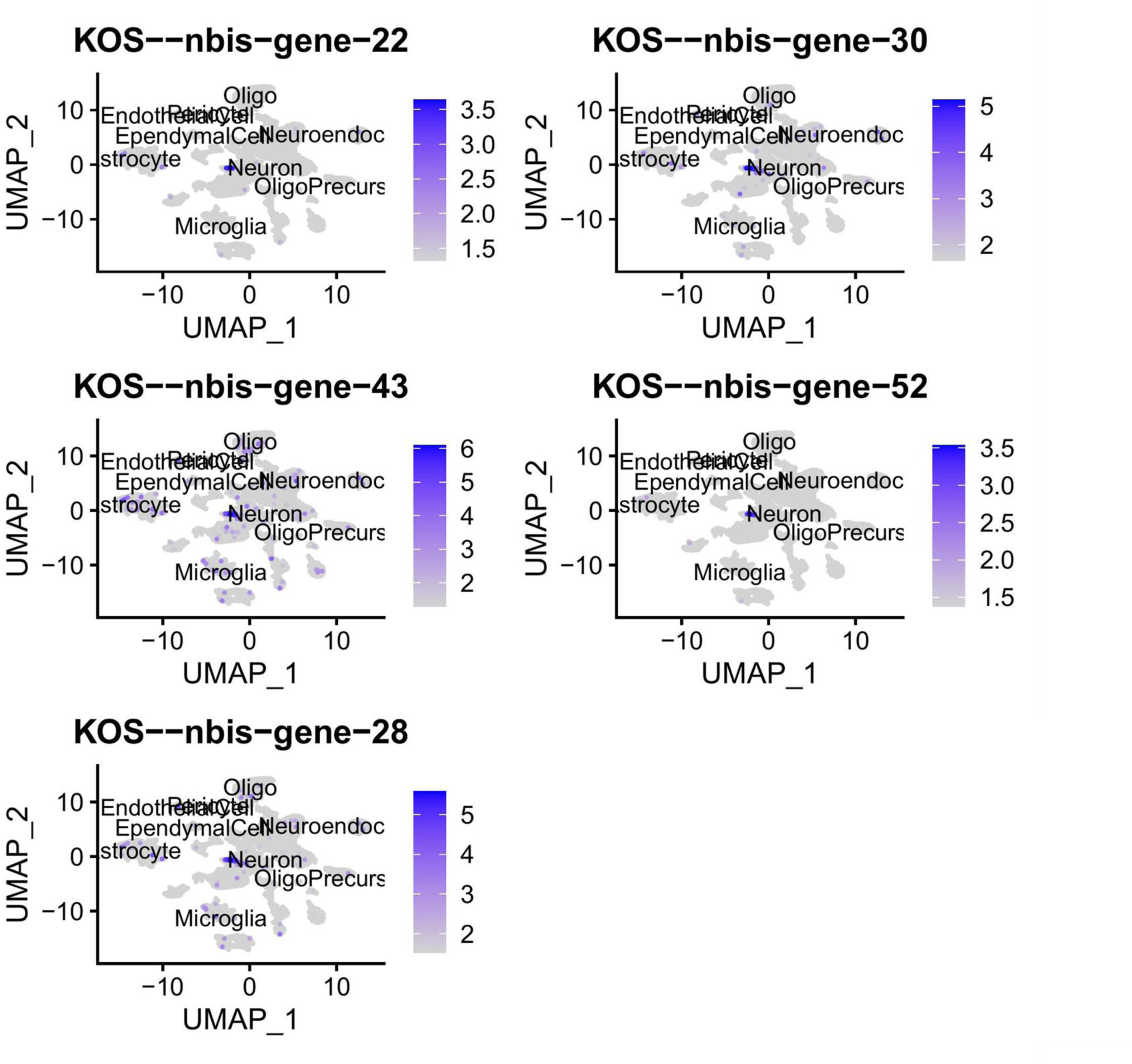
Active viral infection were observed in neuronal and glial cells. UMAP plot of scRNAseq data showing the expression of HSV infected mouse genes in different cell-types

**Fig 8c.**
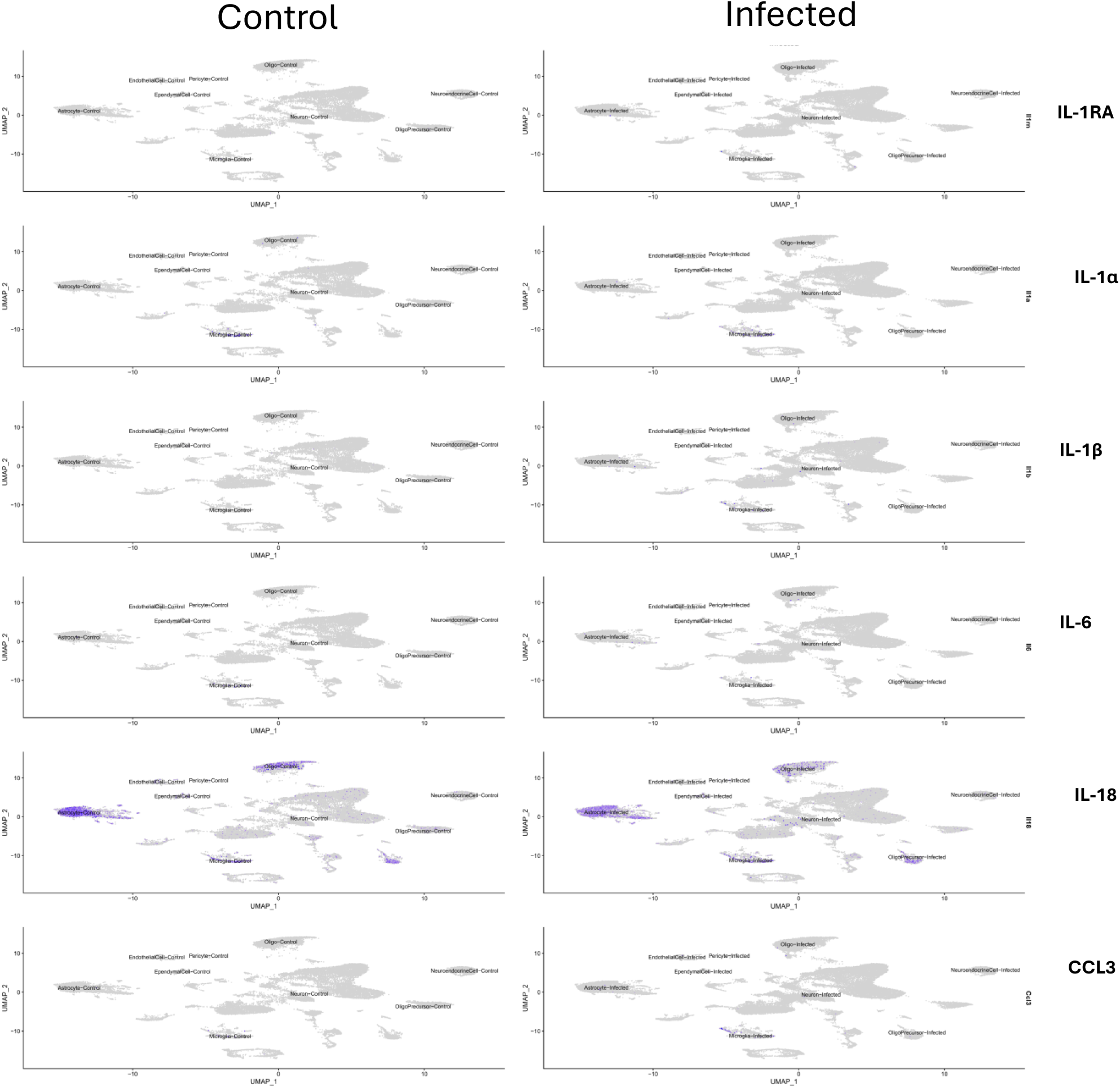
Mediators were abundantly expressed in neuronal and glial cells. Abundance of 6 selected gene expression levels in both control (left) and infected (right) mouse cell types. Colour intensity represents the level of gene expression of each mediator in the specific cell type. Genes selected were: Il1a, Il1b, Il1ra, Il16, Il118, and ccl3.

**Fig 8d.**
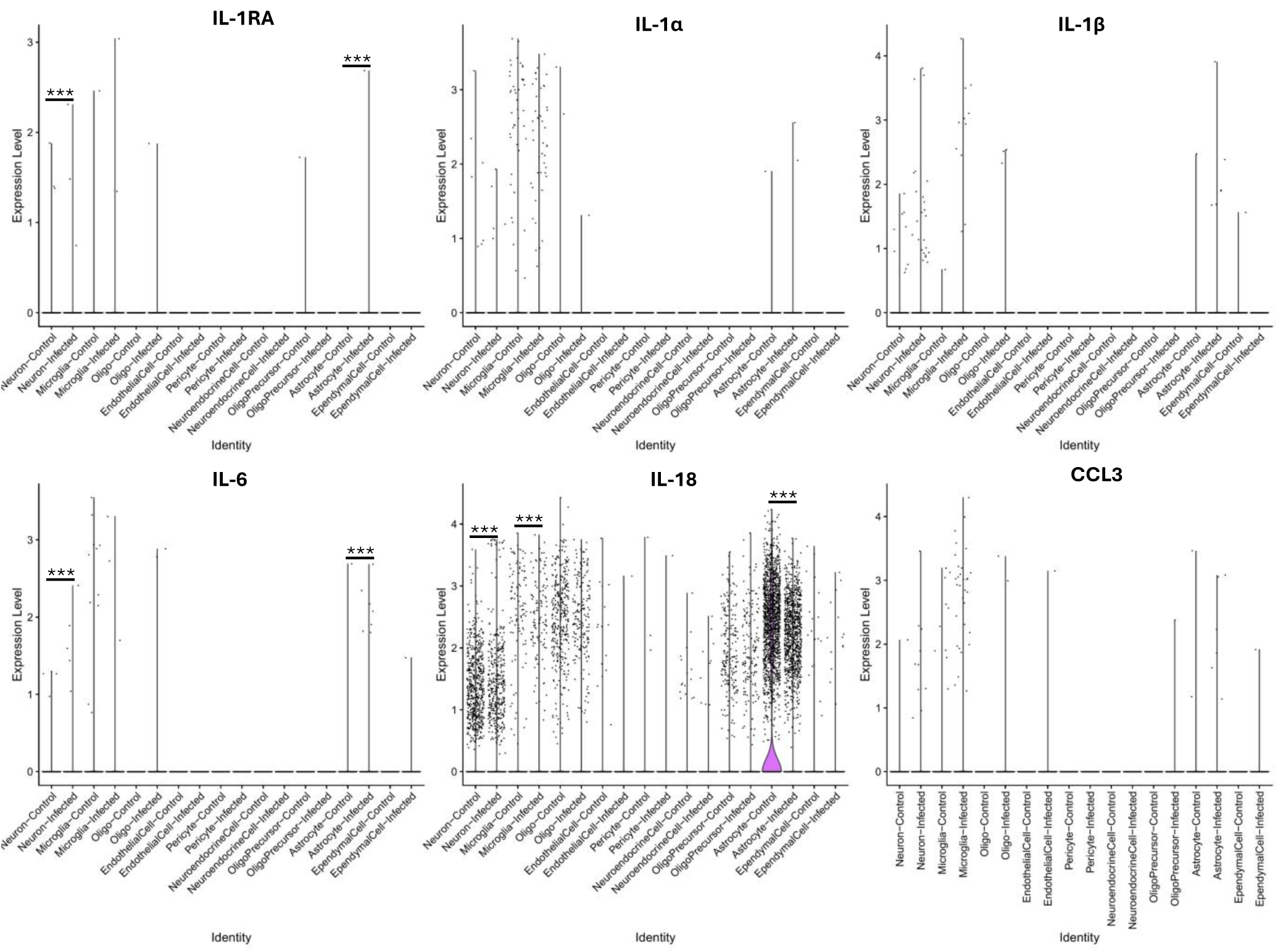
HSV infection induced neuronal, microglial and astrocytic expression of the interleukin-1 cytokines and IL-6. Expression levels of selected mediators in uninfected and infected mouse brain cell types. P values were determined by Bonferroni corrected non-parametric Wilcoxon rank sum test. Bar plots show mean expression levels. Each dot represents an expression in one cell

## Discussion

In adult HSV encephalitis patients, neuroinflammation, with increased cytokines of the IL-1 superfamily and IL-6, are associated with biomarker and radiological evidence of neuroglial injury, clinical severity and poor outcomes. Specifically, worse clinical outcomes were associated with neuroinflammation and neuroglial injury during acute infection, as shown by increased CSF and serum levels of the neuroglial biomarkers GFAP, Tau, and UCH-L1 along with increased CSF and serum levels of IL-1RA, CSF IL-18 and serum IL-6. These changes were associated with increased volume of cerebral oedema on MRI, suggesting that heightened CNS-derived IL-1 cytokines, and an IL-6 mediated systemic inflammatory response may drive neuropathology in HSV encephalitis. Moreover, poor outcomes following HSV encephalitis are more likely driven by associated neuroinflammation and neuroglial injury mediated by IL-6 and IL-1 superfamily cytokines. CSF and serum levels of GFAP, NfL and UCHL-1 are elevated in several neurological disorders, including HSV encephalitis, SARS-CoV-2, and traumatic brain injury ^17–20^. Disease outcomes were not associated with HSV viral load, consistent with mouse models suggesting that the dysregulated excessive immune response, rather than the direct viral cytopathy alone, may mediate the worst outcomes in viral encephalitis^10,11^.

Raised levels of both IL-1RA and IL-18 in acute CSF samples positively correlated with raised levels of the astrocytic protein GFAP and the biomarkers of neuronal injury, Tau protein and UCHL-1, together with increased volume of cerebral oedema on MRI, suggesting that CNS-derived cytokines of the IL-1 superfamily may be driving the associated neuroinflammation and brain injury in HSV encephalitis. Dysregulation of the immune system, leading to excessive production of proinflammatory cytokines, such as IL-1 and IL-6, has been associated with poor outcomes in HSV encephalitis ^7,8,12^, and these were the most consistent mediators associated with neuroinflammation, brain injury, clinical severity and poor outcomes in our study. Our finding that elevated levels of the anti-inflammatory cytokine, IL-1RA, were associated with worse outcomes, neuroglial injury, and neuroinflammation contradicts previous reports and may well reflect the sample timing and the delayed compensation of initial high levels of IL-1α and IL-1β, which have been shown in murine models to rapidly increase immediately following HSV CNS infection ^8,11^.

Dexamethasone neither improved outcomes in our study^9^, nor substantially affected changes in levels of key mediators. Mediator fold-change patterns over time were similar in both the aciclovir-only and aciclovir plus dexamethasone groups, and there was no significant change in mediator levels in matched early and late samples in the aciclovir plus dexamethasone group beyond those seen in the aciclovir-only group. There was also no significant difference in the GCS, LoS, mRS scores, and mediator levels in late samples between the two groups, indicating that dexamethasone did not influence HSV-induced pathology. A previous HSV encephalitis randomised clinical trial, although forced to close early, found no substantial benefits of adjunctive corticosteroid therapy in HSV encephalitis patients ^21^, and a case report described a failed use of corticosteroids to treat an IL-18-driven autoinflammatory condition ^22^. Likewise, intravenous corticosteroids did not improve outcomes in a randomised placebo-controlled trial in adults with traumatic brain injury ^23^. A plausible explanation is that insufficient dexamethasone crossed the blood–brain barrier to directly modulate the cytokines/proteins that we demonstrate here to correlate with HSV-induced neuropathology and outcomes. Especially as we observed that key mediators likely originated within the CNS, and their levels were higher in the CSF than in the blood. Low dexamethasone concentration in brain tissue has been proposed as the possible cause for the failure of effective glucocorticoid treatment in traumatic brain injury^24,25^. Experimental studies suggest that micromolar concentrations of dexamethasone are required in the brain for effective anti-inflammatory and neuroprotective effects ^26^, whereas only traces cross the blood-brain barrier (BBB) when administered systemically under normal BBB conditions ^24,25,27^. It is therefore likely that, in our study, intravenously administered dexamethasone did not reach therapeutic concentrations within the CNS. Instead, its principal clinical anti-inflammatory CNS effect may have been indirect, through stabilisation of BBB integrity and reduced permeability ^28–30^, rather than direct genomic suppression of CNS inflammatory pathways or microglial activation. This suggests that targeted immunomodulatory drugs with improved CNS penetration, alone or in combination with corticosteroids, could prove effective adjunctive treatment alternatives in HSV encephalitis. Anakinra, the pharmacological IL-1R antagonist, for example, has been shown in a murine model of virus-induced neuroinflammation to reduce neurological injury ^11^, which corroborates our data presented here. Also, a combination therapy of anti-IL1 and anti-IL-18 was successfully used to treat an infant with an inherited macrophage activation syndrome associated with elevated serum IL-18, and an acute parainfluenza upper respiratory tract infection ^22^. Thus, drugs that cross the blood-brain barrier and directly reduce IL-1 levels, such as Anakinra and Tadekining α (a recombinant IL-18BP), could be more effective in reducing neuroinflammation and hence, improve outcomes in HSV encephalitis. The effect of immunomodulatory drugs that target HSV-induced IL-1 and IL-6 cytokine function and/or levels is unknown and should be investigated in future human trials, especially considering that dexamethasone treatment was ineffective.

Proinflammatory cytokines, including the IL-1 superfamily and IL-6, were most likely CNS-derived and are highly related to inflammatory pathways with immune regulatory biological functions. In the HSV encephalitis murine model, we observed active acute HSV infection and high expression of IL-1, IL-6 and IL-18 gene transcripts in neuroglial cells - neurons, microglia, and astrocytes, indicating that HSV infection induces neuronal, astrocytic, and microglial expression of the key mediators identified in our HSV encephalitis patient cohort. We therefore suggest that these mediators are likely key drivers of the damaging inflammatory cascade within the CNS.

GO and KEGG analyses indicated that key mediators were related to pathways of other inflammatory syndromes, including influenza A and viral to protein-protein interaction. Our KEGG analysis results and the association between increased CSF IL-18 and IL-1 and increased cerebral oedema volume suggest that the severe neuroinflammation observed in HSV encephalitis may be driven by a highly regulated, lytic inflammatory pathway. The NLRP3 inflammasome is a multi-protein signalling platform that mediates IL-1β and IL-18 production and has been experimentally shown to activate the NLRP3-mediated IL-1β-dependent pathway ^16^, and to play a key role in many inflammatory diseases ^31,32^, including in HSV encephalitis^33^. Interestingly, the NLRP3 inflammasome is sometimes relatively steroid-resistant because it is triggered by intracellular danger signals and cellular stress rather than purely by cytokine transcription, so that once neuronal injury and microglial activation are underway, inflammasome activation may continue despite systemic corticosteroid therapy^34^. Modulating the immune response early during infection, before severe irreversible brain injury occurs, may reduce neuroinflammation and improve clinical outcomes in HSV encephalitis patients ^35^. Human experimental medicine studies evaluating anti-IL-1, IL-6 and IL-18 and the role of the NLRP3 inflammasome are needed to advance mechanistic understanding of the specific inflammatory pathways involved in HSV encephalitis pathology and to determine at what stage of the pathway targeted therapies can reduce neuroglial injury in HSV encephalitis.

Inconsistencies in the timing of sample collection across participants could have introduced variability in the observed differences in mediator levels in matched early and late samples, potentially affecting the comparability of results. In addition, the small sample size of 14 and 20 participants with matched CSF and matched serum samples, respectively, may limit the statistical power of the effect of dexamethasone on mediator levels.

## Conclusion

HSV encephalitis leads to substantial neurological disability and death in many patients despite effective antiviral therapy^1–3^. Avenues to reduce neurological complications and improve outcomes include adjunctive treatment with immunomodulatory drugs to reduce the HSV-induced lytic neurological inflammatory cascade. Our data suggest that CNS-derived IL-1 superfamily and IL-6 cytokines drive HSV-induced neuroinflammation and brain injury, leading to severe clinical disease and worse outcomes, and the levels of these key mediators were unaltered by dexamethasone. Our findings provide evidence of an association between the IL-1 superfamily and IL-6 levels and neuroglial injury, and the ineffectiveness of systemic dexamethasone treatment at significantly modulating these mediators within a cohort of adult HSV encephalitis patients. Immunomodulatory therapy capable of crossing the blood-brain barrier and targeted at the IL-1 superfamily and IL-6 cytokines should be assessed for effectiveness in reducing neuroinflammation and improving outcomes in HSV encephalitis.

## Methods

### Study Participants and Sample Collection

Samples were collected from adults with PCR-confirmed herpes simplex virus (HSV) encephalitis recruited from 53 National Health Service hospitals in the UK, through the DexEnceph study ^36^. The participants were randomised to receive either adjunctive dexamethasone (10 mg 6 hourly for 4 days) plus intravenous aciclovir (10 mg/kg 8 hourly) (treatment group) or aciclovir alone (control group).

From each participant, up to 2.5 mL of CSF was collected in a universal container, and 5 mL of venous blood in a BD Vacutainer Serum Separator Tube (SST). Serum was separated by centrifugation, and both samples were frozen at the sites per local hospital policy. All frozen specimens were couriered in batches to the Infection Neuroscience Laboratory, Clinical Science Centre, University of Liverpool, for analysis. The specimens were stored at −80°C until analysed, and freeze-thaw cycles were minimised. CSF and blood samples were obtained at recruitment, 14 and 30 days after randomisation in the study or when the patient was discharged from the hospital.

### Measurement of Mediators and Brain Injury Biomarkers

A Bio-Plex Pro Human Cytokine 48-plex Assay (cat. no. 12007283, Bio-Rad Laboratories) was used for mediators profiling. Serum samples were diluted fourfold in sample dilution, then 50 μl was pipetted in triplicate into each well. Standards and CSF samples were analysed undiluted, and plates were processed according to the manufacturer’s instructions. Four biomarkers of neuroglial injury: GFAP, NfL, Total Tau protein, and Ubiquitin Carboxyl-terminal Hydrolase L1 (UCHL-1) were assessed using a Quanterix Simoa Human Neurology 4-Plex B Advantage kit (Product no. 103345, Quanterix Corporation). Serum and CSF samples were diluted fourfold and fortyfold, respectively, in sample diluent. Then, 100 μL of samples, standards, and controls were pipetted into each well, and the plates were processed according to the manufacturer’s instructions. Plates were read on the Quanterix Simoa SR-X (ultrasensitive digital ELISA) benchtop reader. We used the manufacturer’s calibrators, provided with the kit, to calculate concentrations.

### Clinical and functional outcome measures

Clinical severity of HSV encephalitis was assessed using the Glasgow coma scale (GCS) score obtained at admission; a score of 15 (out of 15) was defined as normal, and scores of <15 were defined as abnormal or severe clinical disease. Disability and functional outcome assessments were: Modified Rankin Score (mRS); scores less than 3 (out of 6) were defined as good, and scores of 3 and above were poor, the Liverpool outcome score (LOS); a score of 2 (out of 5) was defined as poor, while scores above 2 were defined as good, and the Barthel Index score. mRS is widely used as an outcome measure of disability and dependence of patients following stroke or neurological infections, or non-infectious encephalitis ^37^. LOS assess outcome following Japanese encephalitis, but has been successfully used in adolescents and adults in a previous HSV encephalitis study^38^.

### Magnetic resonance imaging

Magnetic Resonance Imaging (MRI) scans were obtained from patients’ standard medical care records between admission and 7 days post-randomisation. A second scan was done two weeks after randomisation. Volumes of total, cytotoxic and vasogenic oedema and temporal lobe volumes were assessed as previously described and established for patients with HSV encephalitis ^36^. Temporal lobe volume was calculated as a percentage of intracranial volume.

### Herpes simplex virus encephalitis mouse model

C57BL/6 female mice aged 8 -12 weeks were intracranially infected with 20 μl of 10^4^ PFU HSV-1-KOS ^39^ or sham-infected with sterile PBS. Mice were euthanised with CO_2_ three days post-infection, and death was confirmed by transection of the great arteries. Post-mortem, 0.5 cm of the cerebral cortex was sampled into RNAlater and immediately processed for single-nuclei isolation.

#### Single Nuclei Isolation

Samples were delicately dounced homogenised in 2ml of ice-cold Nuclei EZ lysis buffer (Sigma, #EZ PREP NUC-101), transferred into a different tube containing an additional 2ml of ice-cold Nuclei EZ lysis buffer and incubated on ice for 5min. Samples were then filtered through a 30μm cell strainer (MACS Smart Strainer 130-098-458) and centrifuged at 500g for 5min at 4°C. The supernatant was discarded, and the pellet was resuspended in fresh 4mL of Nuclei EZ lysis buffer. This process was repeated three times. After the last centrifugation, the pellets were resuspended in 300μL of NSB (1xPBS, 5% BSA, 0.25% Glycerol and 40U/μL RNAse inhibitor from #Clontech Takara 2313A/B) and filtered through a 40 μm Flowmi Cell Strainer (H13680-0040), followed by filtration through a 10 μm cell strainer. The single nuclei suspension was stained with 1 μg/ml DAPI (Fisher, Cat # D1306) and DAPI+ events were sorted. The concentration of single nuclei was adjusted to 1000 single nuclei per μL and immediately loaded on the 10X Chromium controller instrument (10X Genomics) for sequencing.

#### Single-cell RNA sequencing

Cell suspensions were loaded along with reverse transcriptase reagents, 3′ gel beads, and emulsification oil onto separate channels of a 10X Single Cell B Chip, which was loaded into the 10X Chromium instrument to generate emulsions. Emulsions were transferred to polymerase chain reaction (PCR) strip tubes for immediate processing and reverse transcription. Library preparation was performed according to the manufacturer’s recommendations. Expression libraries were generated using Chromium Single Cell Next GEM 3′V3.1 chemistry (10X Genomics PN-1000268). DNA and library quality were evaluated using an Agilent 2100 Bioanalyzer, and concentration was quantified using the Qubit dsDNA high-sensitivity reagents (Thermo Fisher Scientific). Gene expression libraries were sequenced on the Illumina NextSeq 500/550 High Output Kit v2.5 (75 Cycles) (Illumina #20024906).

### Ethical approval

Informed consent was provided for all participants before recruited into the study. All ethical approvals were in place before the start of the clinical trial. Ethics approval was obtained from the UK National Research Ethics Committee (reference number: 15/NW/0545).

### Statistical Analysis

R statistical software version 4.5.0 (R Core Team 2025) was used for the statistical analysis. Categorical and numerical variables were compared between groups using the Chi-Squared and independent samples Wilcoxon rank-sum (Mann-Whitney) tests, respectively. Numerical variables between matched groups were compared using the Wilcoxon signed-rank test. Correlations were assessed with either Spearman rho or Kendall tau rank correlation tests. A *P* value of 0.05 was considered statistically significant. Graphs were generated using the ‘ggplot2’ package (Wickham, 2016), and statistical annotations were added using the ‘ggpubr’ package (Kassambara, 2024).

To avoid undetectable levels or missing data bias, only mediators detected in CSF or serum specimens from ≥80% of the samples were analysed. Also, mediators with low variability, for which over 90% of the measured concentrations were at or below the limit of quantification, were excluded from all analyses. To minimise any influence of sample storage, concentrations of each mediator were median-centred and normalised for each participant. Therefore, the concentration of each mediator was expressed and analysed as a value relative to the median concentration of all of the mediators in that participant’s sample, as described previously ^40,41^.

Differential gene expression analysis, for murine data, was done using the Seurat R package (version 5.2.0). Gene expression between groups was compared with the Wilcoxon rank sum test using the FindMarkers() function. The *P* values were corrected by the Bonferroni method to control for multiple comparisons.

#### Kyoto Encyclopedia of Genes and Genomes (KEGG) Protein Enrichment Analysis

The UniProt IDs of mediators that were significantly different between participants with GCS scores <15 and a GCS score of 15 were obtained from https://uniprot.org. Before the enrichment analysis, UniProt names were first converted to ‘Entrez IDs’ using the ‘org.Hs.eg.db’ annotation package (Carlson, 2024). KEGG pathway enrichment analysis was done using the enrichKEGG() function from the clusterProfiler R package (Yu et al., 2012). Parameters of the analysis were a *P* value cutoff of 0.05 and the Benjamini-Hochberg method for multiple testing correction. Pathway classifications from the KEGG map search results were ranked by the highest number of mapped candidates and exported in the KGML format. Key pathway diagrams were generated and visualised using the ‘pathview’ package (Luo & Brouwer, 2020). It overlaid the log2 fold-change values of each mediator, providing a visual representation of their regulation within biological contexts.

#### STRING protein-protein interaction analysis

STRING v12, accessed at https://string-db.org was used for network analysis, with proteins entered as a list using their UniProt accessions and searched against the Homo sapiens database. In all cases, data were filtered so that only interactions identified in experimental, or database-level evidence were included at medium confidence, and any disconnected nodes were removed for viewing. KMEANS clustering was used to identify clusters of proteins with shared functions, the number of clusters set was determined empirically for each network. Networks generated in STRING were exported as a tab-separated values (.tsv) file and imported into Cytoscape v3.9.1 for editing and PowerPoint to customize the overall network presentation.

#### Gene Ontology - functional enrichment analysis

DAVID Bioinformatics Resources v2024q4 was used to understand the functional enrichment of proteins identified in this study, using a background database of Homo sapiens. Data was filtered so that GO terms: Molecular Function (MF) and Biological Process (BP) were maintained. *P* values were adjusted using the Benjamini-Hochberg method. Data visualization was performed using a custom R script and labelling amended in InkScape v1.2.

## Acknowledgements

The authors thank the patients and their carers who participated in this study, and staff in all the study hospitals.

---

**Extended Figure 1.**
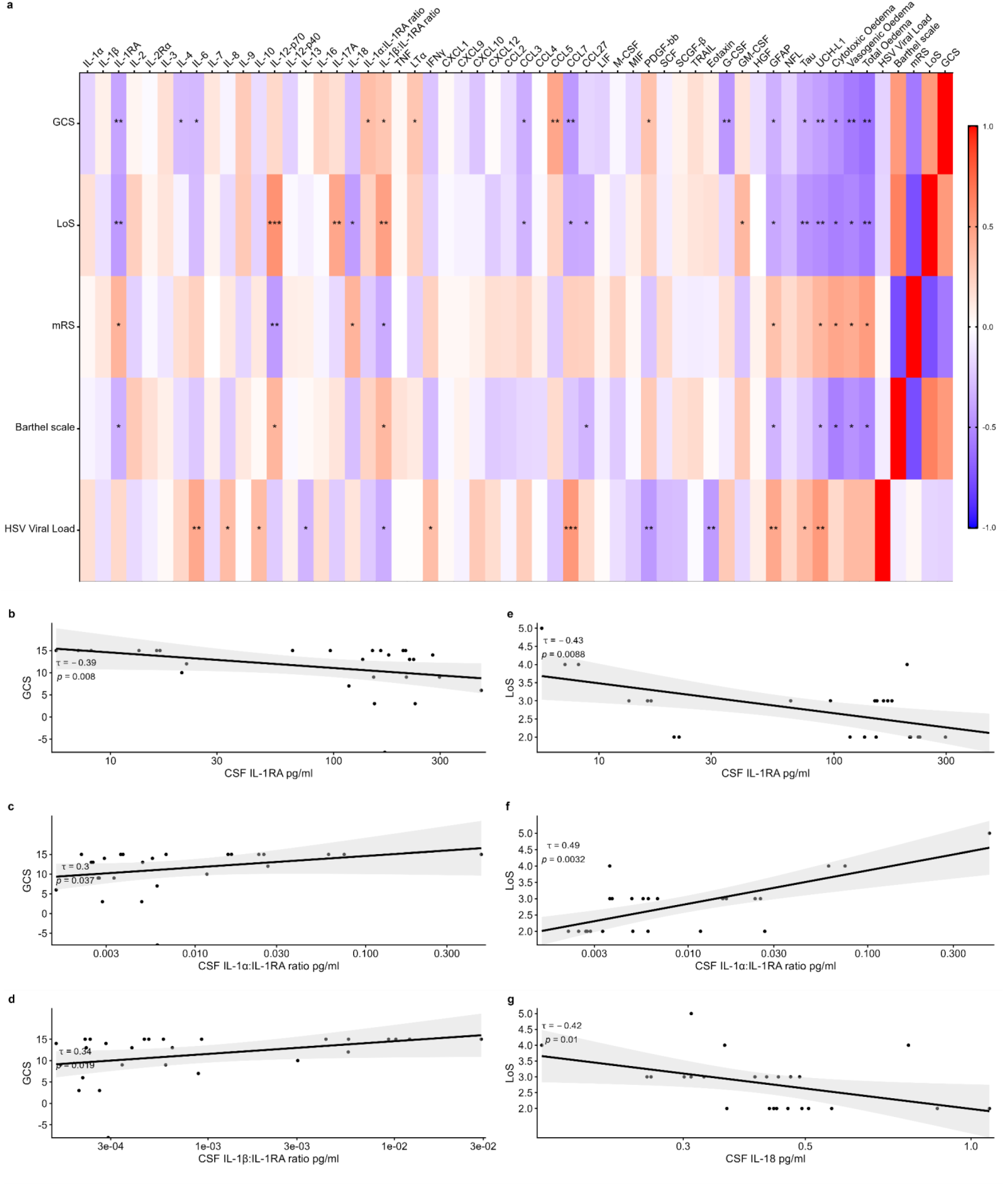
Association between clinical severity, outcome scores and biomarkers in CSF. **a)** Kendall’s correlation matrix of mediators, HSV viral load, GFAP, NfL, T-Tau, UCHL1, GCS, LOS, mRS, and the Barthel Scale. **b – d)** correlation between GCS scores and (b) IL-1RA, (c) IL-1α:IL-1RA ratio, and (d) IL-1β:IL-1RA ratio ; **e-g)** association between LOS and (e) IL-1RA, (f) IL-1α:IL-1RA ratio, and (g) IL-18. * = *P* < 0.05, ** = *P* < 0.01, and *** = *P* < 0.001

**Extended Figure 2.**
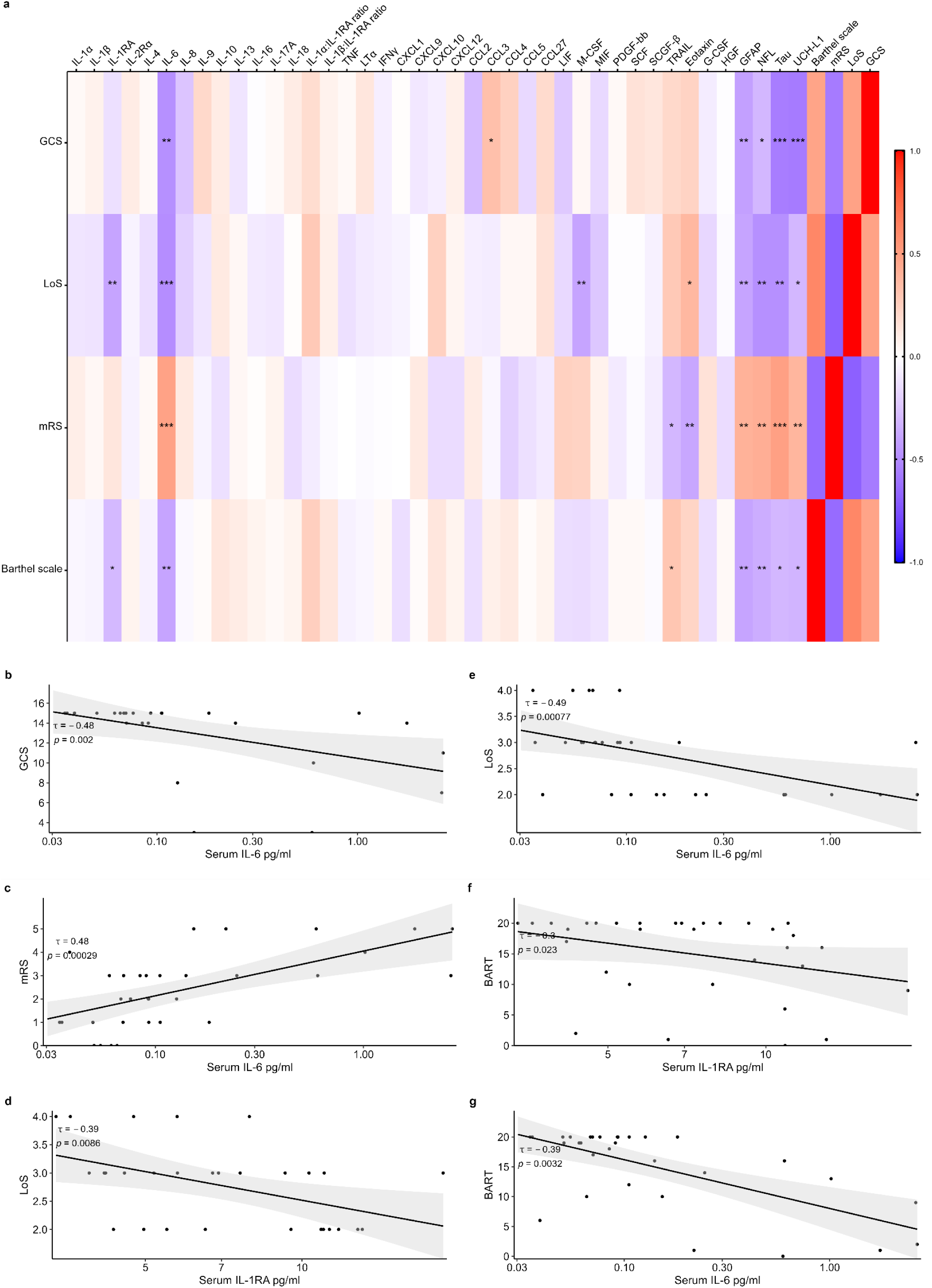
Association between clinical severity, outcome scores and biomarkers in serum. **a)** Kendall’s correlation matrix of mediators, HSV viral load, GFAP, NFL, T-Tau, UCHL1, GCS, LoS, mRS, and the Barthel Scale. **b-g)** correlation plots between (b) GCS and IL-6, (c) mRS and IL-6; (d) Los and IL-1β:IL-1RA ratio; (e) LoS and IL-6; (f) Barth scale score and IL-1RA, and (g) Barth scale score and IL-6. * = *P* < 0.05, ** = *P* < 0.01, and *** = *P* < 0.001

**Extended Figure 3.**
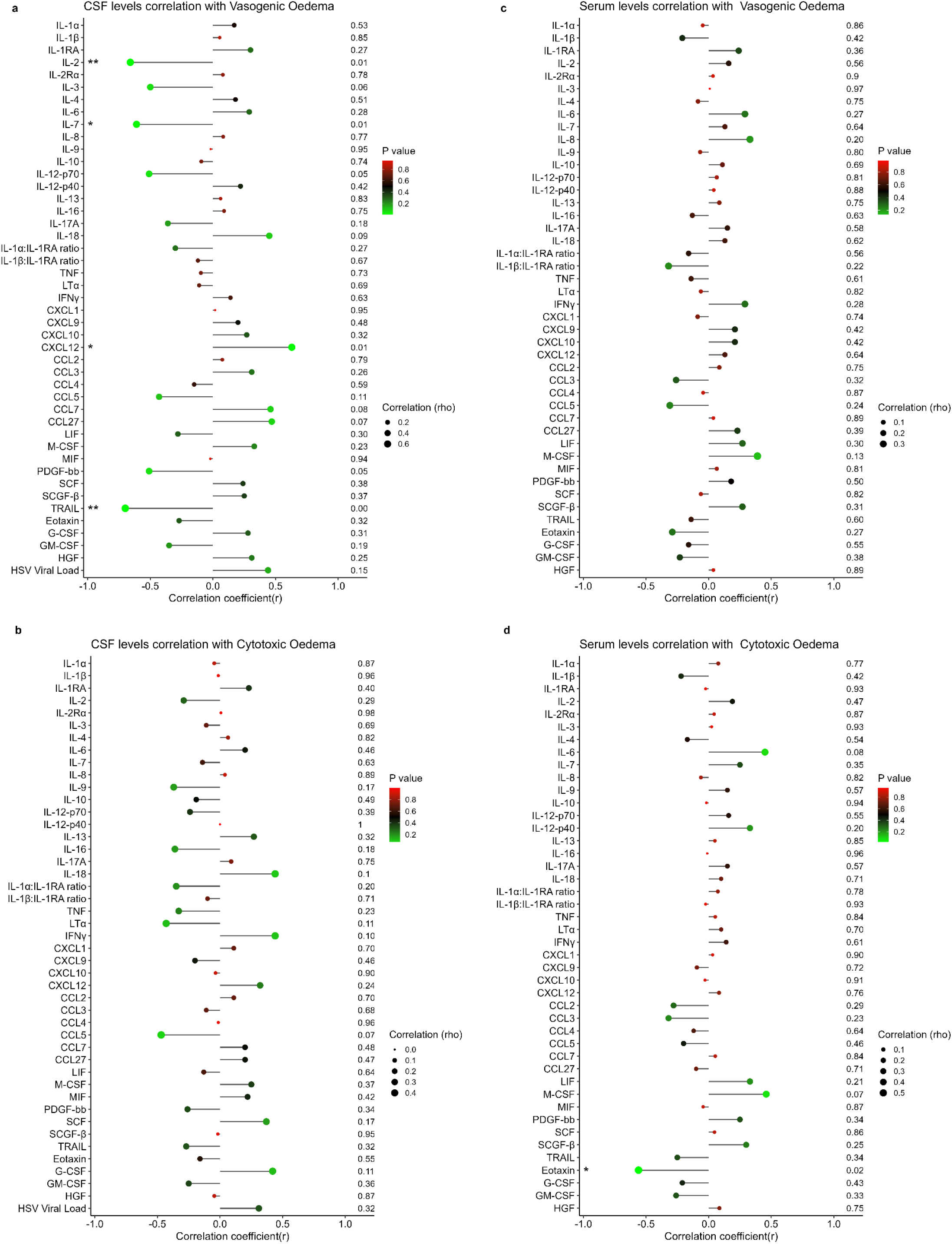
Correlation between acute mediator levels and MRI scans. a-d, Forest plot of Spearman correlations between mediators and MRI scan. a, correlation between CSF mediator levels and Vasogenic oedema; b, correlation between CSF mediator levels and Cytotoxic oedema; c, correlation between Serum mediator levels and Vasogenic oedema; d, correlation between Serum mediator levels and Cytotoxic oedema. Dot size corresponds to the Spearman correlation coefficient, while colour represents the P value. * = P < 0.05, ** = P < 0.01, and *** = P < 0.001

**Extended Figure 4.**
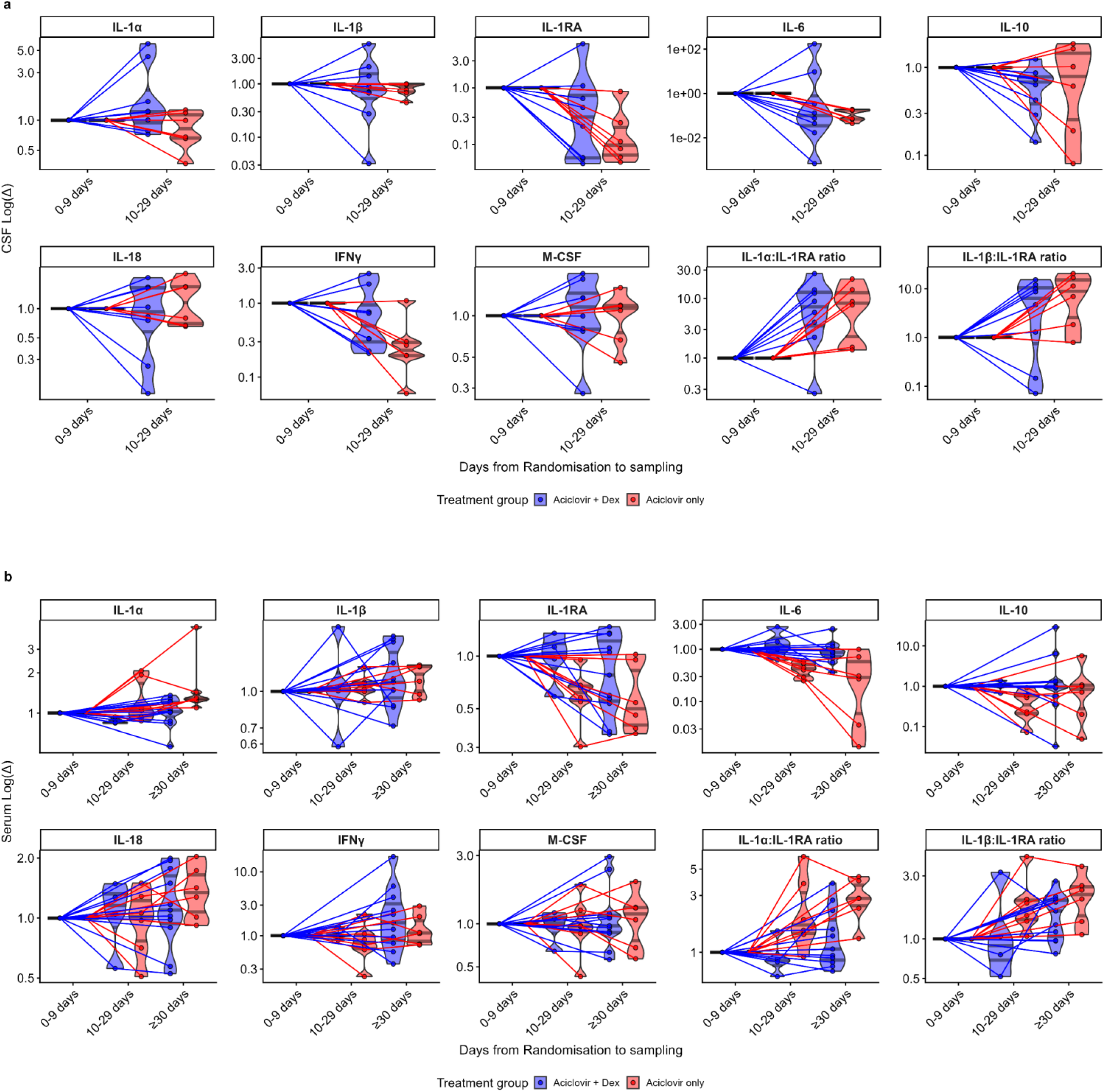
Fold changes in mediator levels were unrelated to adjunct dexamethasone treatment. Transient log fold changes in mediator levels, stratified by treatment group in **(a) CSF: n=14** and **(b) Serum: n=20**. Each line represents matched longitudinal samples from an individual participant collected at 0–9, 10-29 and ≥30 days post-randomisation. Levels at 0-9 days were normalised to 1, and subsequent levels are expressed relative to it. Only serum was collected at ≥30 days post-randomisation. The violin plots show the median and interquartile levels; each data point represents one patient’s clinical sample. For participants with multiple samples within a given time window, only the earliest sample was included in the analysis

**Extended Figure 5.**
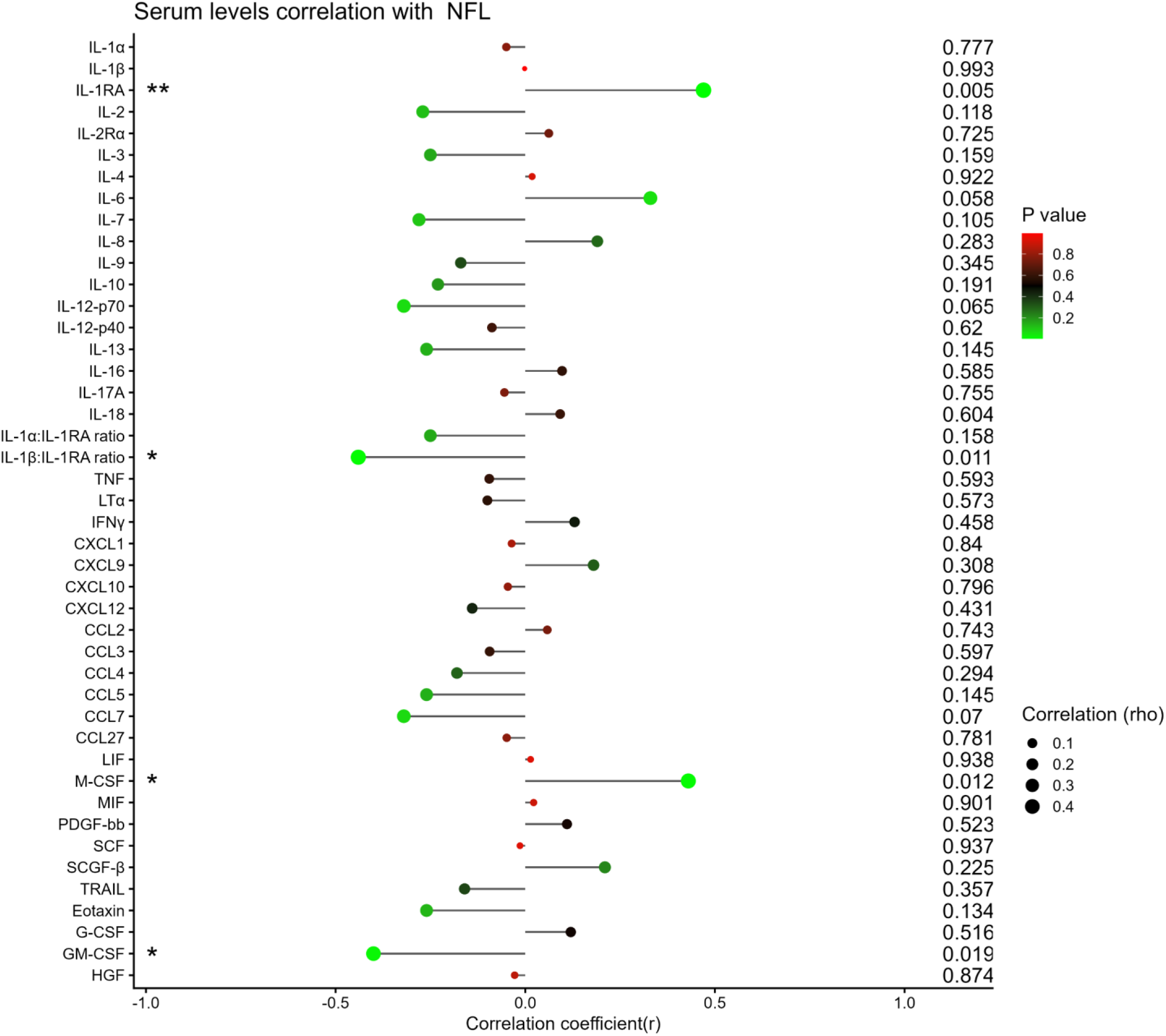
Association between blood levels of mediators and Neurofilament Light chain (NfL). Forest plot of Spearman correlation between concentration of mediators and NfL in serum. Dot size corresponds to the Spearman correlation coefficient, while colour represents the **P** value. * = P < 0.05, ** = P < 0.01, and *** = P < 0.001

**Extended Figure 6.**
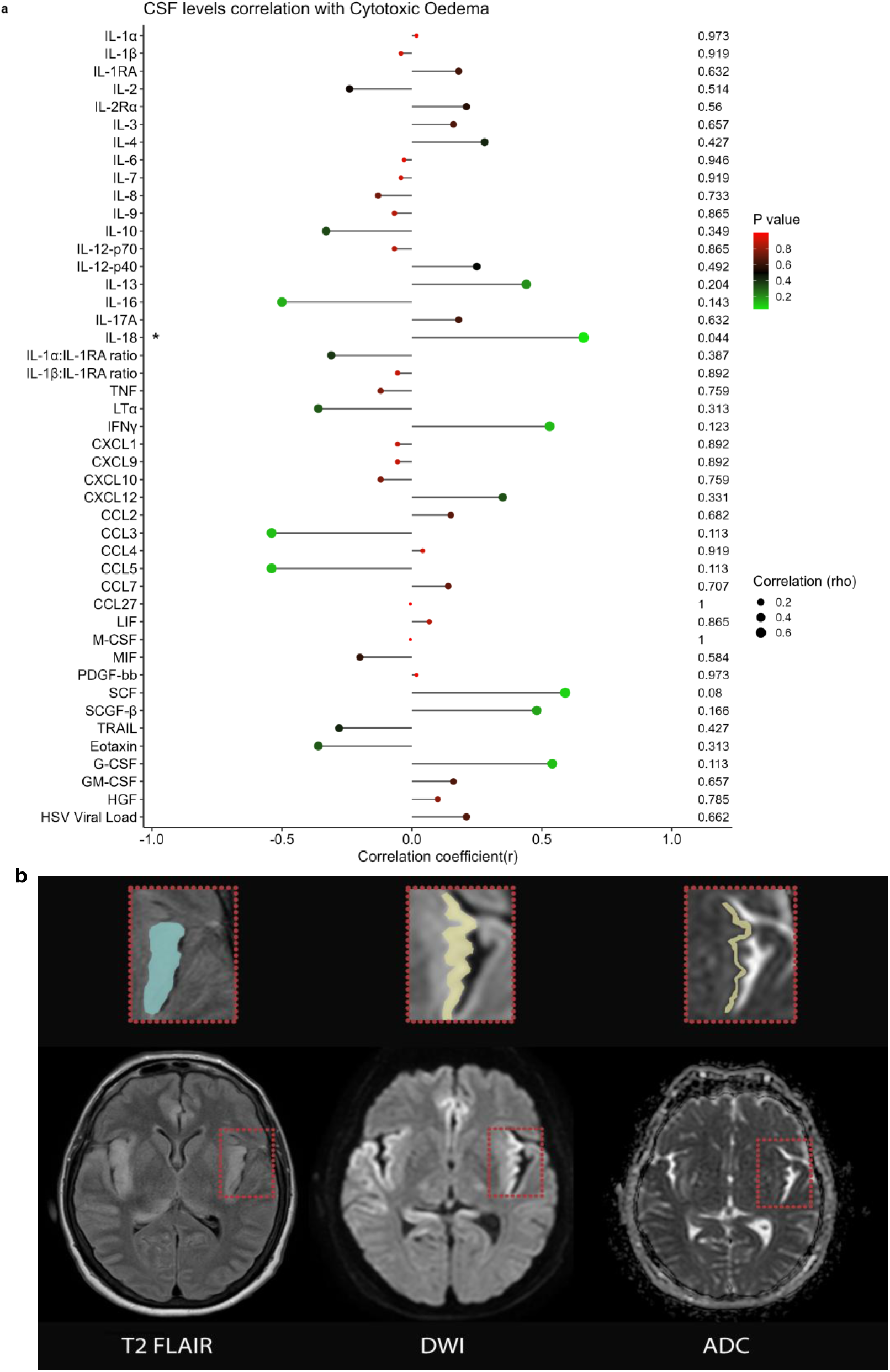
Association between CSF levels of mediators and MRI. **a,** Forest plot of Spearman correlation between CSF concentration of mediators and Cytotoxic oedema for 10 participants with an MRI scan and CSF sample collected within 3 days. Dot size corresponds to the Spearman correlation coefficient, while colour represents the *P* value. **b**, representative MRI sequences showing cytotoxic and vasogenic oedema in an adult HSV encephalitis patient. Axial T2 FLAIR, diffusion-weighted imaging (DWI) and apparent diffusion coefficient (ADC) maps. Red dashed boxes indicate the region of interest enlarged in the upper panels. A representative T2 FLAIR hypersensitivity is highlighted in cyan, reflecting total oedema burden encompassing both cytotoxic and vasogenic components. Cytotoxic oedema is identified by the combination of DWI hypersensitivity and corresponding ADC hypointensity, highlighted in yellow, indicating restricted diffusion consistent with cellular swelling. Regions of T2 FLAIR hyperintensity without restricted diffusion on DWI/ADC represent vasogenic oedema, attributed to blood—brain barrier disruption and increased extracellular water

**Extended Figure 7.**
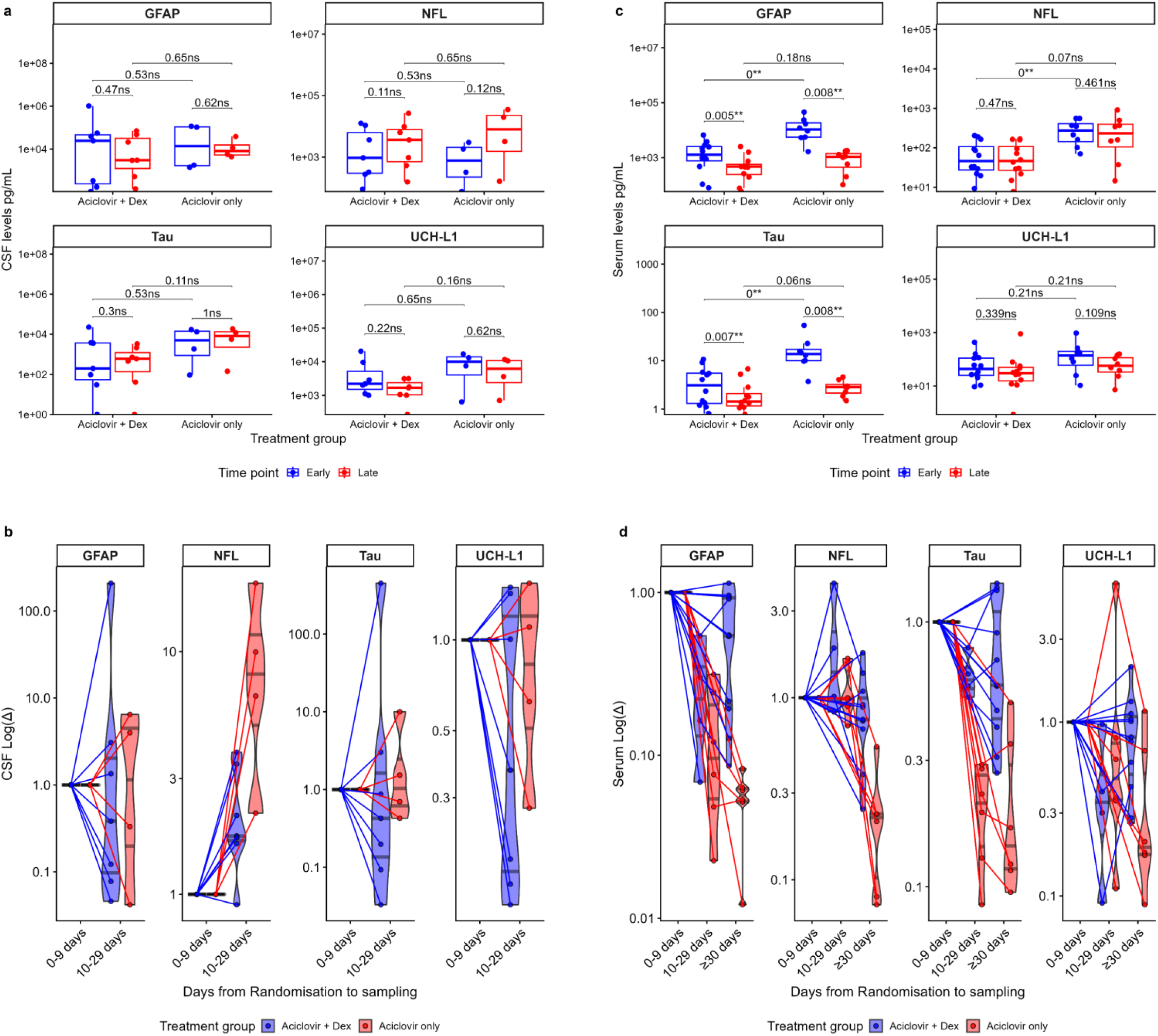
Changes in CSF and blood levels of brain injury biomarkers were unrelated to adjunct dexamethasone treatment. a,c. Boxplots comparing mediator levels in matched early and late clinical samples, stratified by treatment group in **(a) CSF**: 11 participants and (**c) Serum**: 20 participants. Matched samples within each group were compared using the Wilcoxon signed-rank test, and unpaired comparisons between groups using the Mann-Whitney U test. The boxplots show the median and interquartile values; each data point represents one patient’s clinical sample. Samples collected within 9 days of randomisation were defined as early, and samples collected on or after day 10 post-randomisation as late. b,d. Transient log fold changes in mediator levels, stratified by treatment group in **(b) CSF**: n=14 and **(d) Serum**: n=20. Each line represents matched longitudinal samples from an individual participant collected at 0–9, 10-29 and ≥30 days post-randomisation. Levels at 0-9 days were normalised to 1, and subsequent levels are expressed relative to it. Only serum was collected at ≥30 days post-randomisation. The boxplots and violin plots show the median and interquartile levels; each data point represents one patient’s clinical sample. For participants with multiple samples within a given time window, only the earliest sample was included in the analysis

**Extended Figure 8.**
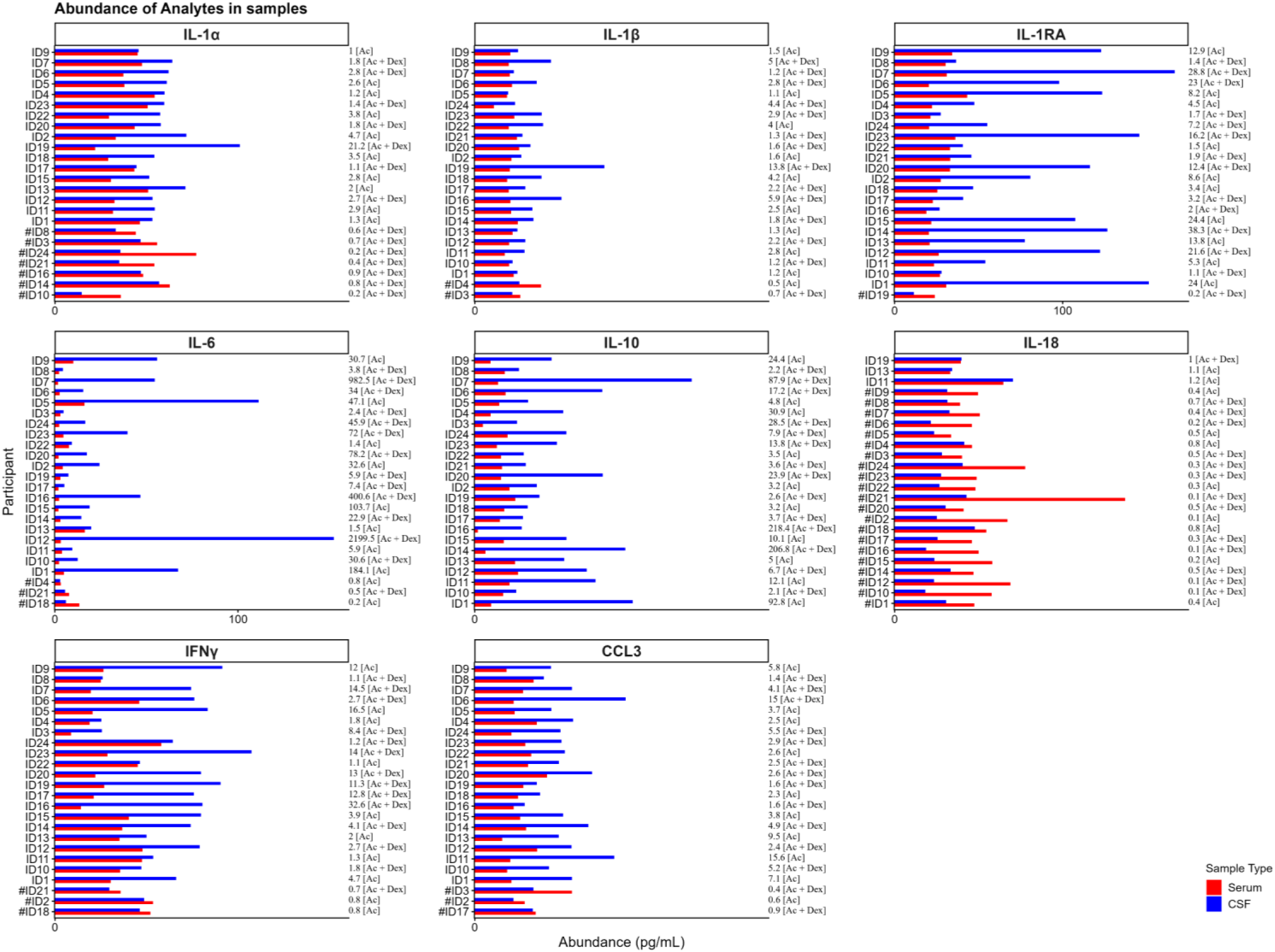
Key mediator levels were higher in CSF compared to blood. Analyte abundance plots comparing concentrations of mediators in 24 patient-matched CSF and serum samples collected within 0-9 days after randomisation. Each analyte represents median-centred and normalised values. The ratios of CSF/Serum levels and the treatment groups are displayed on the right. Ac = Aciclovir only and Ac + Dex = Aciclovir and Dexamethasone; # denotes participants with higher levels in serum than CSF

