## Supplementary data for "Central nervous system–derived inflammasome cytokines drive neuroglial injury and cerebral oedema in viral encephalitis, despite corticosteroid therapy"

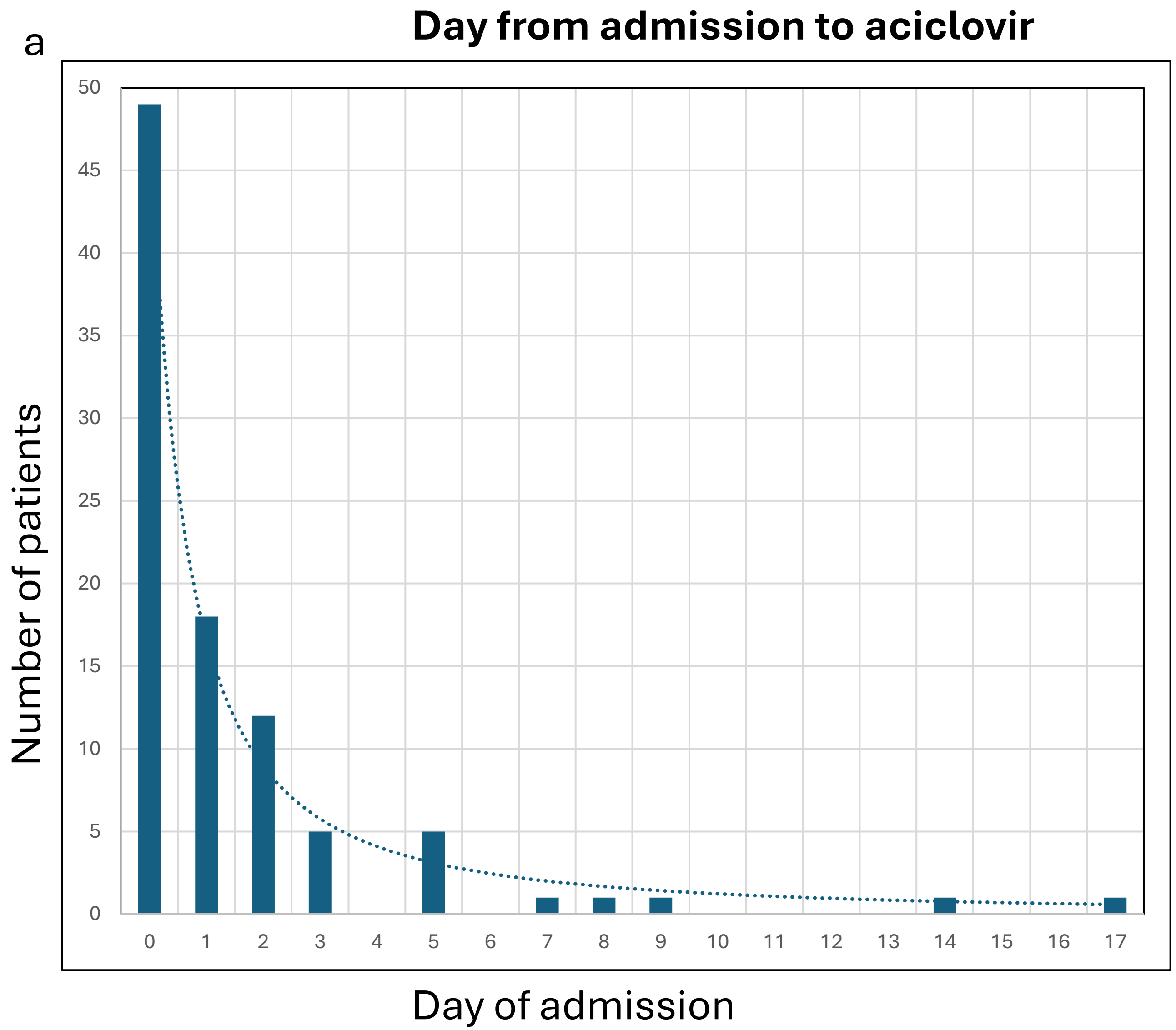


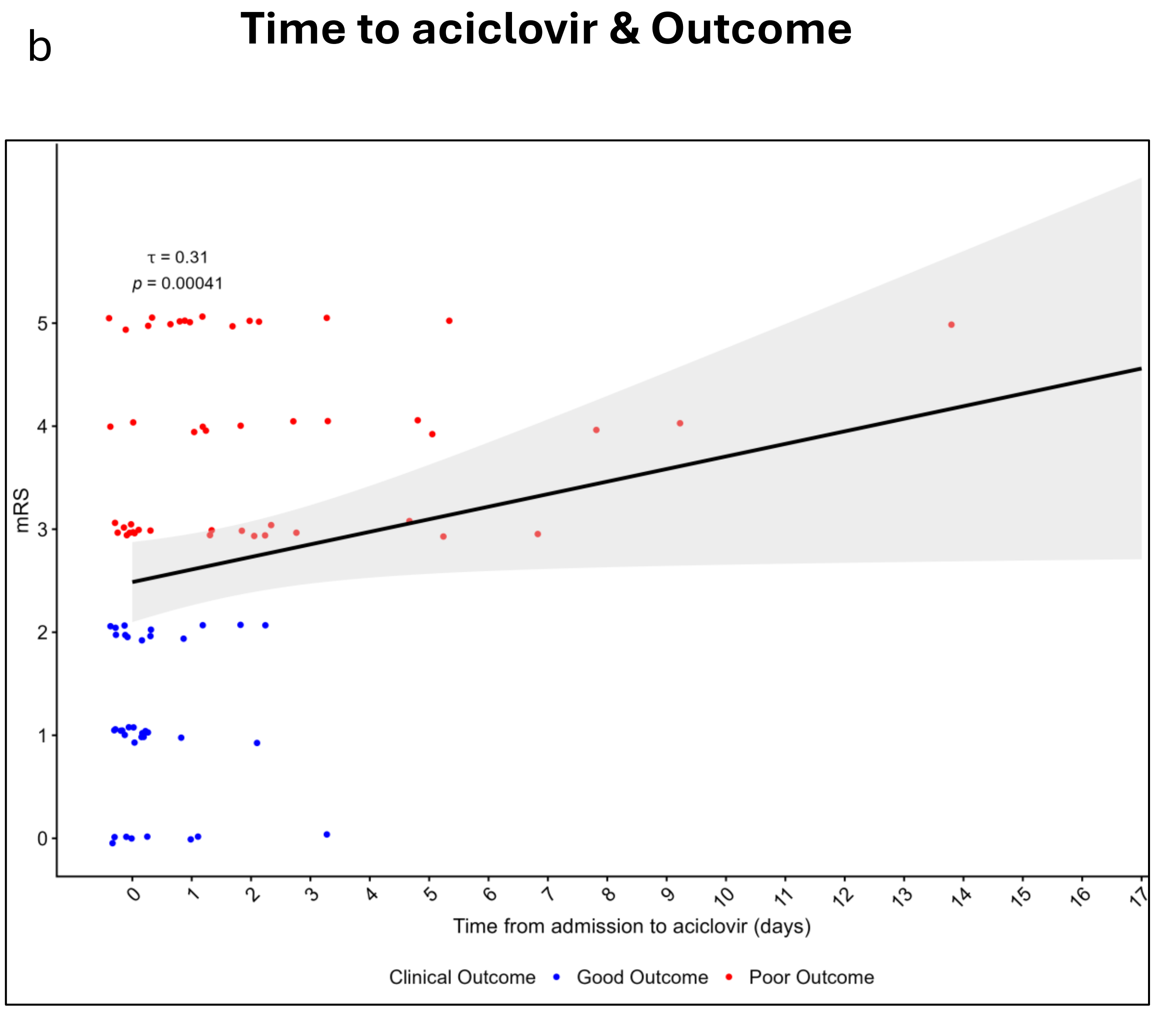


**Supplementary Figure 1a,b. Early aciclovir initiation was associated with improved outcomes.**
a) Bar plots of the number of patients per day from admission to aciclovir initiation; Each bar plot shows the number of patients. (b) Kendall's correlation plot between mRS and time from admission to aciclovir initiation; each data point represents one patient.

C

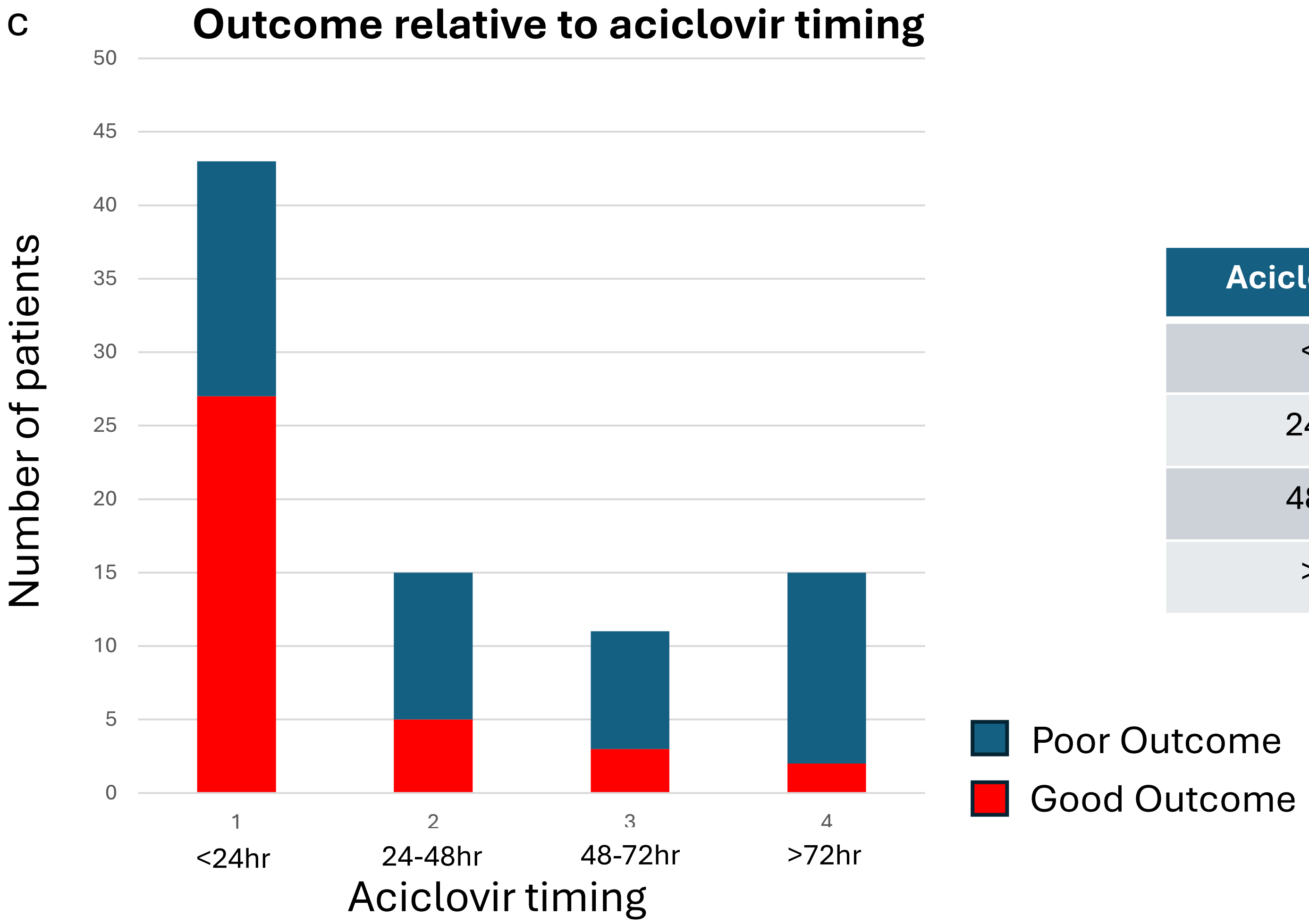


| Aciclovir Timing | Good Outcome (%) | Poor Outcome (%) |
|---|---|---|
| <24hrs | 62.8 | 37.2 |
| 24-48hrs | 33.3 | 66.7 |
| 48-72hrs | 27.3 | 72.7 |
| >72hrs | 13.3 | 86.7 |

**Supplementary Figure 1c.** Bar plots of the number of patients per day from admission to aciclovir initiation stratified by clinical outcome.

| mRS | Timing of Aciclovir | | | | Odd Ratio Good Outcome (95%CI) | | |
|---|---|---|---|---|---|---|---|
| | **<24hrs** | **24-48hrs** | **48-72hrs** | **>72hrs** | **<24 vs 24-28** | **<24 vs 48-72** | **<24 vs >72** |
| Good | 27 | 5 | 3 | 2 | **3.38 (1.01-12.53)** | **4.5 (1.12-22.89)** | **10.97 (2.60-76.18)** |
| Poor | 16 | 10 | 8 | 13 | | | |

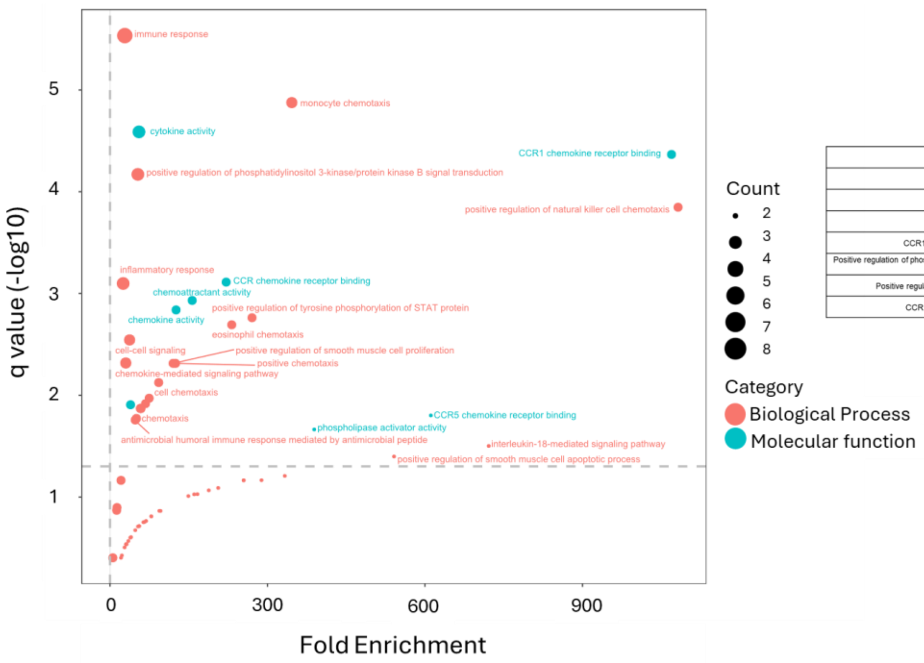


| Category | Proteins in category | Bonferroni q-value |
|---|---|---|
| Immune response | CCL3, CCL5, CSF3, IFNG, IL1RN, IL12A, IL18 | 2.92E-06 |
| Cytokine activity | CSF3, IFNG, IL1RN, IL12A, IL18 | 2.58E-05 |
| Monocyte chemotaxis | CCL3, CCL5, CCL7, PDGFB | 2.67E-05 |
| CCR1 chemokine receptor binding | CCL3, CCL5, CCL7 | 0.000086 |
| Positive regulation of phosphatidylinositol 3-kinase/protein kinase B signal transduction | CCL3, CCL5, CSF3, IL18, PDGFB | 0.000203 |
| Positive regulation of natural killer cell chemotaxis | CCL3, CCL5, CCL7 | 0.000566 |
| CCR chemokine receptor binding | CCL3, CCL5, CCL7 | 0.00232 |

**Supplementary Figure 2. Gene ontology functional analyses highlight immune regulatory functions of key mediators.**
GO enrichment analysis of the 9 significant proteins was performed in DAVID functional annotation analysis. P-values were adjusted for multiple hypothesis testing using the Benjamini-Hochberg correction and plotted against fold enrichment. The horizontal line represents an adjusted p-value of 0.05. Points are sized according to the number of proteins within each enriched term and colored according to the enrichment category.

**Supplementary Table 1:** Acute Blood and CSF concentrations of all measured mediators, brain injury biomarkers and HSV viral load grouped by clinical severity and outcome scores

### CSF

| | GCS | | | mRS (30days) | | | LoS (30days) | | | Barthel (30 days) | | |
|---|---|---|---|---|---|---|---|---|---|---|---|---|
| Biomarker | Normal (GCS = 15) n = 11 | Abnormal (GCS < 15) n = 12 | p value | Good Outcome (mRS<3) n = 11 | Poor Outcome (mRS>2) n = 11 | p value | Good recovery (LoS > 2) n = 13 | Poor Recovery (LoS =2) n = 9 | p value | Independent (score =20) n = 12 | High dependency (score <20) n = 10 | p value |
| IL-1α | 0.58 ( 0.43 - 0.68 ) | 0.68 ( 0.57 - 0.73 ) | NS | 0.63 ( 0.53 - 1 ) | 0.62 ( 0.44 - 0.7 ) | NS | 0.63 ( 0.47 - 0.93 ) | 0.62 ( 0.48 - 0.69 ) | NS | 0.6 ( 0.43 - 0.81 ) | 0.66 ( 0.53 - 0.72 ) | NS |
| *IL-1β* | 0.07 ( 0.06 - 0.09 ) | 0.06 ( 0.05 - 0.09 ) | NS | 0.06 ( 0.05 - 0.07 ) | 0.08 ( 0.05 - 0.1 ) | NS | 0.06 ( 0.05 - 0.08 ) | 0.08 ( 0.05 - 0.1 ) | NS | 0.07 ( 0.05 - 0.09 ) | 0.06 ( 0.05 - 0.1 ) | NS |
| IL-1RA | 41.06 ( 12.11 - 152.66 ) | 194.26 ( 139.07 - 231.13 ) | 0 | 81.03 ( 12.11 - 164.33 ) | 151.84 ( 93.28 - 216.11 ) | NS | 81.03 ( 14.09 - 160.07 ) | 180.72 ( 121.55 - 224.37 ) | 0.03 | 81.03 ( 15.43 - 164.33 ) | 151.84 ( 93.28 - 216.11 ) | NS |
| IL-2 | **0.27 ( 0.16 - 0.32 )** | 0.18 ( 0.14 - 0.26 ) | NS | 0.29 ( 0.19 - 0.35 ) | 0.18 ( 0.14 - 0.23 ) | NS | **0.29 ( 0.21 - 0.37 )** | 0.16 ( 0.14 - 0.19 ) | 0.03 | 0.29 ( 0.25 - 0.35 ) | 0.16 ( 0.13 - 0.2 ) | 0.01 |
| IL-2Rα | 0.85 ( 0.76 - 1.41 ) | 0.96 ( 0.76 - 1.66 ) | NS | 0.99 ( 0.79 - 1.41 ) | 0.96 ( 0.68 - 2.09 ) | NS | 1.06 ( 0.81 - 1.63 ) | 0.91 ( 0.6 - 1.64 ) | NS | **1.17 ( 0.82 - 2.01 )** | 0.91 ( 0.55 - 1.22 ) | NS |
| IL-3 | 0.06 ( 0.04 - 0.08 ) | 0.04 ( 0.04 - 0.07 ) | NS | 0.06 ( 0.04 - 0.08 ) | 0.04 ( 0.04 - 0.06 ) | NS | 0.06 ( 0.04 - 0.08 ) | 0.05 ( 0.04 - 0.07 ) | NS | 0.06 ( 0.04 - 0.08 ) | 0.04 ( 0.03 - 0.06 ) | NS |
| IL-4 | 0.07 ( 0.06 - 0.09 ) | 0.08 ( 0.07 - 0.12 ) | NS | 0.08 ( 0.06 - 0.12 ) | 0.08 ( 0.07 - 0.09 ) | NS | 0.08 ( 0.06 - 0.12 ) | 0.08 ( 0.07 - 0.09 ) | NS | 0.08 ( 0.06 - 0.12 ) | 0.08 ( 0.07 - 0.09 ) | NS |
| IL-6 | 3.77 ( 0.36 - 30.16 ) | 28.08 ( 5.76 - 44.96 ) | NS | 5.65 ( 0.36 - 23.98 ) | 28.8 ( 2.68 - 64.96 ) | NS | 5.65 ( 0.6 - 28.24 ) | 28.8 ( 6.2 - 44.96 ) | NS | 5.65 ( 0.98 - 23.98 ) | 28.8 ( 2.37 - 64.96 ) | NS |
| IL-7 | 1.17 ( 0.78 - 1.47 ) | 1 ( 0.55 - 1.2 ) | NS | 1.09 ( 0.78 - 1.39 ) | 1.17 ( 0.56 - 1.41 ) | NS | 1.09 ( 0.89 - 1.33 ) | 1.17 ( 0.55 - 1.43 ) | NS | 1.17 ( 0.97 - 1.43 ) | 1.06 ( 0.47 - 1.41 ) | NS |
| IL-8 | 9.08 ( 5.02 - 34.93 ) | 14.92 ( 5.79 - 23.11 ) | NS | 6.01 ( 3.07 - 16.43 ) | 11.29 ( 4.49 - 26.21 ) | NS | 6.01 ( 3.06 - 12.5 ) | 19.04 ( 5.91 - 26.86 ) | NS | 5.74 ( 3.04 - 16.43 ) | 12.58 ( 7.82 - 26.21 ) | NS |
| IL-9 | 0.81 ( 0.56 - 1.15 ) | 0.62 ( 0.5 - 0.73 ) | NS | 0.74 ( 0.54 - 1.15 ) | 0.66 ( 0.5 - 0.75 ) | NS | 0.71 ( 0.56 - 1.1 ) | 0.66 ( 0.46 - 0.73 ) | NS | 0.7 ( 0.54 - 1.04 ) | 0.66 ( 0.5 - 0.76 ) | NS |
| IL-10 | 0.36 ( 0.26 - 1.16 ) | 0.78 ( 0.44 - 1.75 ) | NS | 0.75 ( 0.26 - 1.37 ) | 0.47 ( 0.34 - 0.85 ) | NS | 0.41 ( 0.28 - 1.29 ) | 0.6 ( 0.36 - 1.01 ) | NS | 0.75 ( 0.3 - 1.37 ) | 0.47 ( 0.33 - 0.85 ) | NS |
| IL-12-p70- | 0.16 ( 0.12 - 0.23 ) | 0.07 ( 0.06 - 0.13 ) | 0.01 | 0.16 ( 0.11 - 0.23 ) | 0.09 ( 0.06 - 0.12 ) | 0.02 | 0.19 ( 0.12 - 0.23 ) | 0.07 ( 0.06 - 0.11 ) | 0 | 0.16 ( 0.11 - 0.23 ) | 0.09 ( 0.06 - 0.12 ) | 0.02 |
| IL-12-p40- | 2.92 ( 2.53 - 3.35 ) | 3.98 ( 3.39 - 4.86 ) | NS | 3.11 ( 2.53 - 4.87 ) | 3.59 ( 2.88 - 4.85 ) | NS | 3.11 ( 2.7 - 4.64 ) | 3.59 ( 2.83 - 4.86 ) | NS | 3.11 ( 2.83 - 4.87 ) | 3.59 ( 2.65 - 4.85 ) | NS |
| IL-13 | 0.11 ( 0.09 - 0.25 ) | 0.13 ( 0.07 - 0.17 ) | NS | 0.11 ( 0.08 - 0.14 ) | 0.15 ( 0.09 - 0.18 ) | NS | 0.11 ( 0.08 - 0.19 ) | 0.15 ( 0.1 - 0.17 ) | NS | 0.11 ( 0.09 - 0.25 ) | 0.13 ( 0.08 - 0.17 ) | NS |
| IL-16 | 2.31 ( 0.84 - 3.19 ) | 1.21 ( 1.01 - 1.81 ) | NS | 2.25 ( 0.82 - 3.19 ) | 1.39 ( 1.01 - 1.95 ) | NS | 1.85 ( 0.98 - 2.85 ) | 1.46 ( 1.02 - 2 ) | NS | 2.25 ( 0.99 - 3.19 ) | 1.39 ( 1.01 - 1.95 ) | NS |
| IL-17A | 0.28 ( 0.25 - 0.34 ) | 0.23 ( 0.18 - 0.28 ) | NS | 0.28 ( 0.24 - 0.32 ) | 0.24 ( 0.18 - 0.3 ) | NS | 0.29 ( 0.24 - 0.33 ) | 0.22 ( 0.17 - 0.26 ) | 0.03 | 0.28 ( 0.24 - 0.32 ) | 0.24 ( 0.18 - 0.3 ) | NS |
| IL-18 | 0.38 ( 0.31 - 0.45 ) | 0.5 ( 0.44 - 0.62 ) | 0.01 | 0.34 ( 0.31 - 0.44 ) | 0.46 ( 0.44 - 0.52 ) | 0.02 | 0.34 ( 0.3 - 0.44 ) | 0.48 ( 0.44 - 0.55 ) | 0.01 | 0.38 ( 0.31 - 0.45 ) | 0.45 ( 0.41 - 0.52 ) | NS |
| TNF | 0.96 ( 0.79 - 1.04 ) | 1.02 ( 0.84 - 1.07 ) | NS | 0.98 ( 0.9 - 1.22 ) | 0.91 ( 0.78 - 1.04 ) | NS | 0.97 ( 0.86 - 1.15 ) | 0.94 ( 0.77 - 1.06 ) | NS | 0.98 ( 0.95 - 1.22 ) | 0.85 ( 0.75 - 1.04 ) | NS |
| LTα | 1.28 ( 1.01 - 1.81 ) | 0.98 ( 0.86 - 1.22 ) | NS | 1.22 ( 0.88 - 1.81 ) | 1.06 ( 0.83 - 1.2 ) | NS | 1.15 ( 0.92 - 1.68 ) | 1.01 ( 0.82 - 1.22 ) | NS | 1.13 ( 0.88 - 1.49 ) | 1.06 ( 0.83 - 1.24 ) | NS |
| IFNγ | 0.87 ( 0.52 - 1.54 ) | 1.69 ( 1.15 - 2.03 ) | 0.05 | 1.18 ( 0.55 - 1.54 ) | 1.62 ( 0.88 - 1.98 ) | NS | 1.18 ( 0.55 - 1.57 ) | 1.62 ( 1.01 - 2.03 ) | NS | 1.18 ( 0.52 - 1.54 ) | 1.62 ( 0.9 - 1.98 ) | NS |
| CXCL1 | 12.69 ( 4.84 - 28.85 ) | 4.39 ( 0.92 - 14.96 ) | NS | 11.21 ( 2.59 - 23.53 ) | 11.74 ( 1.77 - 22.75 ) | NS | 12.69 ( 3.5 - 25.97 ) | 8.76 ( 1.62 - 17.3 ) | NS | 12.69 ( 2.59 - 23.53 ) | 8.76 ( 1.77 - 22.75 ) | NS |
| CXCL9 | 61.4 ( 10 - 117.44 ) | 46.03 ( 33.43 - 204.92 ) | NS | 102.59 ( 10 - 158.52 ) | 38.36 ( 28.6 - 141.18 ) | NS | 101.49 ( 16.04 - 144.37 ) | 34.49 ( 23.36 - 185.45 ) | NS | 112.52 ( 24.4 - 158.52 ) | 34.49 ( 18.05 - 95.39 ) | NS |
| CXCL10 | 510.3 ( 264.81 - 1115.38 ) | 857.07 ( 384.56 - 1546.01 ) | NS | 848.67 ( 329.5 - 1177.02 ) | 478.3 ( 345.6 - 1557.2 ) | NS | 848.67 ( 305.76 - 1295.31 ) | 478.3 ( 370.72 - 1333.45 ) | NS | 848.67 ( 264.81 - 1177.02 ) | 554.86 ( 410.27 - 1557.2 ) | NS |
| CXCL12 | 17.06 ( 4.64 - 24.49 ) | 18.4 ( 12.05 - 26.47 ) | NS | 17.06 ( 4.64 - 24.49 ) | 19.55 ( 10.69 - 27.46 ) | NS | 14.61 ( 3.98 - 23.55 ) | 24.13 ( 12.95 - 28.61 ) | NS | 15.09 ( 3.58 - 24.49 ) | 19.81 ( 12.33 - 27.46 ) | NS |
| CCL2 | 11.02 ( 5.3 - 23.52 ) | 17.1 ( 7.77 - 40.19 ) | NS | 7.91 ( 4.45 - 14.14 ) | 19.52 ( 8.93 - 29.93 ) | NS | 10.46 ( 4.86 - 22.03 ) | 17.1 ( 7.77 - 34.34 ) | NS | 7.91 ( 4.45 - 14.14 ) | 19.52 ( 8.93 - 29.03 ) | NS |
| CCL3 | 0.16 ( 0.09 - 0.26 ) | 0.26 ( 0.22 - 0.34 ) | 0.02 | 0.19 ( 0.11 - 0.29 ) | 0.25 ( 0.19 - 0.29 ) | NS | 0.19 ( 0.1 - 0.28 ) | 0.26 ( 0.22 - 0.32 ) | NS | 0.19 ( 0.11 - 0.29 ) | 0.25 ( 0.19 - 0.29 ) | NS |
| CCL4 | 0.95 ( 0.74 - 1.73 ) | 0.93 ( 0.82 - 1.5 ) | NS | 0.87 ( 0.81 - 1.39 ) | 0.89 ( 0.8 - 1.3 ) | NS | 0.84 ( 0.76 - 1.25 ) | 0.93 ( 0.83 - 1.52 ) | NS | 0.84 ( 0.74 - 1.08 ) | 0.93 ( 0.82 - 1.65 ) | NS |
| CCL5 | 1.43 ( 1.08 - 1.91 ) | 0.69 ( 0.49 - 1 ) | 0 | 1.23 ( 0.97 - 1.87 ) | 0.78 ( 0.46 - 1.54 ) | NS | 1.23 ( 0.94 - 1.71 ) | 0.76 ( 0.45 - 1.67 ) | NS | 1.23 ( 0.97 - 1.55 ) | 0.78 ( 0.46 - 1.79 ) | NS |
| CCL7 | 0.3 ( 0.17 - 0.62 ) | 1.19 ( 0.68 - 2.03 ) | 0 | 0.38 ( 0.21 - 0.81 ) | 0.75 ( 0.39 - 1.61 ) | NS | 0.38 ( 0.2 - 0.72 ) | 1.06 ( 0.48 - 1.84 ) | NS | 0.38 ( 0.21 - 0.81 ) | 0.75 ( 0.39 - 1.61 ) | NS |
| CCL27 | 0.6 ( 0.42 - 0.76 ) | 0.67 ( 0.49 - 0.94 ) | NS | 0.53 ( 0.42 - 0.74 ) | 0.73 ( 0.53 - 0.99 ) | NS | 0.53 ( 0.42 - 0.73 ) | 0.77 ( 0.56 - 1 ) | NS | 0.53 ( 0.42 - 0.74 ) | 0.73 ( 0.53 - 0.99 ) | NS |
| LIF | 1.77 ( 1.5 - 2 ) | 1.94 ( 1.74 - 2.28 ) | NS | 1.83 ( 1.7 - 2.21 ) | 1.83 ( 1.62 - 2.07 ) | NS | 1.83 ( 1.6 - 2.15 ) | 1.83 ( 1.66 - 2.09 ) | NS | 1.78 ( 1.54 - 1.99 ) | 1.94 ( 1.67 - 2.14 ) | NS |
| M-CSF | 1.74 ( 0.91 - 3.2 ) | 1.36 ( 1.25 - 2.09 ) | NS | 1.22 ( 0.83 - 1.89 ) | 1.62 ( 1.28 - 3.25 ) | NS | 1.36 ( 0.86 - 2.05 ) | 1.6 ( 1.25 - 2.83 ) | NS | 1.22 ( 0.83 - 1.89 ) | 1.62 ( 1.28 - 3.12 ) | NS |
| MIF | 38.91 ( 26.15 - 58.68 ) | 44.28 ( 32.92 - 62.12 ) | NS | 38.91 ( 33.23 - 52.75 ) | 43.94 ( 29.83 - 65.54 ) | NS | 40.61 ( 35.58 - 61.21 ) | 40.28 ( 27.84 - 55.13 ) | NS | 38.91 ( 33.23 - 52.75 ) | 43.94 ( 29.83 - 67.22 ) | NS |
| PDGF-bb | 3.84 ( 3.26 - 4.44 ) | 2.93 ( 1.83 - 3.69 ) | 0.04 | 3.76 ( 3.09 - 4.44 ) | 3.03 ( 2.14 - 3.98 ) | NS | 3.76 ( 2.84 - 4.25 ) | 3.03 ( 2.38 - 3.97 ) | NS | 3.84 ( 3.09 - 4.44 ) | 3.03 ( 2.14 - 3.59 ) | NS |
| SCF | 1.39 ( 0.96 - 1.91 ) | 1.52 ( 1.26 - 2.04 ) | NS | 1.31 ( 0.82 - 1.59 ) | 1.52 ( 1.08 - 2.16 ) | NS | 1.31 ( 0.86 - 1.77 ) | 1.52 ( 1.18 - 2.04 ) | NS | 1.31 ( 0.82 - 1.59 ) | 1.52 ( 1.08 - 2.07 ) | NS |
| SCGF-β | 899.36 ( 566.38 - 1156.48 ) | 664.2 ( 409.91 - 1216.18 ) | NS | 698.32 ( 585.4 - 1108.36 ) | 711.8 ( 391.38 - 1215.72 ) | NS | 698.32 ( 549.76 - 1137.49 ) | 711.8 ( 427.55 - 1216.18 ) | NS | 867.39 ( 585.4 - 1156.48 ) | 696.16 ( 391.38 - 941.38 ) | NS |
| TRAIL | 0.31 ( 0.26 - 0.64 ) | 0.25 ( 0.22 - 0.29 ) | NS | 0.29 ( 0.23 - 0.4 ) | 0.27 ( 0.22 - 0.36 ) | NS | 0.29 ( 0.23 - 0.43 ) | 0.27 ( 0.22 - 0.35 ) | NS | 0.31 ( 0.26 - 0.56 ) | 0.25 ( 0.21 - 0.33 ) | NS |
| Eotaxin | 0.15 ( 0.12 - 0.22 ) | 0.14 ( 0.11 - 0.16 ) | NS | 0.16 ( 0.13 - 0.22 ) | 0.14 ( 0.11 - 0.15 ) | NS | 0.15 ( 0.13 - 0.2 ) | 0.14 ( 0.11 - 0.15 ) | NS | 0.15 ( 0.12 - 0.22 ) | 0.14 ( 0.12 - 0.16 ) | NS |
| G-CSF | 6.94 ( 5.52 - 12.12 ) | 12.76 ( 11.44 - 14.87 ) | 0.01 | 9.66 ( 6.01 - 14.34 ) | 12.47 ( 7.92 - 14.57 ) | NS | 9.66 ( 5.78 - 14.43 ) | 12.47 ( 8.96 - 14.2 ) | NS | 7.13 ( 5.52 - 13.08 ) | 12.76 ( 10.39 - 14.57 ) | NS |
| GM-CSF | 0.19 ( 0.16 - 0.2 ) | 0.11 ( 0.09 - 0.19 ) | NS | 0.19 ( 0.16 - 0.21 ) | 0.14 ( 0.1 - 0.19 ) | NS | 0.19 ( 0.17 - 0.22 ) | 0.11 ( 0.1 - 0.17 ) | 0.02 | 0.19 ( 0.18 - 0.21 ) | 0.11 ( 0.09 - 0.18 ) | 0.01 |
| HGF | 19.16 ( 17.14 - 21.66 ) | 25.22 ( 12.3 - 36.44 ) | NS | 21.09 ( 17.46 - 26.04 ) | 18.8 ( 12.08 - 35.42 ) | NS | 21.09 ( 17.28 - 27.32 ) | 18.8 ( 11.89 - 34.6 ) | NS | 20.97 ( 17.14 - 26.04 ) | 20.61 ( 12.08 - 35.42 ) | NS |
| GFAP | 2212.75 ( 308.76 - 26719.24 ) | 100820.04 ( 35518.95 - 132151.05 ) | 0 | 11384.5 ( 850.9 - 34023.86 ) | 82328.51 ( 2497.47 - 107603.76 ) | NS | 11384.5 ( 336.84 - 38914.41 ) | 96418.17 ( 8069.58 - 113034.46 ) | NS | 11384.5 ( 308.76 - 34023.86 ) | 82328.51 ( 2497.47 - 107603.76 ) | NS |
| NFL | 590.94 ( 185.59 - 1230.12 ) | 2121.94 ( 1234.55 - 6219.28 ) | 0.03 | 945.51 ( 209.47 - 2352.06 ) | 1543.34 ( 630.01 - 3889.9 ) | NS | 945.51 ( 279.25 - 1292.65 ) | 1821.36 ( 803.22 - 5646.53 ) | NS | 945.51 ( 303.13 - 2352.06 ) | 1543.34 ( 630.01 - 3889.9 ) | NS |
| Tau | 224.33 ( 131.52 - 2341.86 ) | 5693.43 ( 1328.2 - 16275 ) | 0.01 | 703.29 ( 131.52 - 3052.9 ) | 2990.51 ( 1008.37 - 13423.41 ) | NS | 476.19 ( 96.82 - 2540.57 ) | 5693.43 ( 1588.23 - 13885.17 ) | 0.01 | 703.29 ( 181.51 - 3052.9 ) | 2990.51 ( 1008.37 - 13423.41 ) | NS |
| UCH-L1 | 2037.71 ( 1251.16 - 4088.23 ) | 10792.85 ( 7557.86 - 16372.52 ) | 0 | 2219.05 ( 1483.54 - 5990.15 ) | 8869.98 ( 4841.41 - 15324.34 ) | 0.02 | 2219.05 ( 1379.13 - 7689.89 ) | 10384.78 ( 5676.82 - 16372.52 ) | 0.01 | 2219.05 ( 1483.54 - 5990.15 ) | 8869.98 ( 4841.41 - 15324.34 ) | 0.03 |
| HSV viral load (copies/ml) | 6093.04 ( 435.45 - 85194.02 ) | 30375.86 ( 17246.9 - 100419.04 ) | NS | 6713.36 ( 1741.8 - 92279.94 ) | 28215.88 ( 12149.67 - 59332.34 ) | NS | 6713.36 ( 870.9 - 88258.2 ) | 37948.2 ( 17237.16 - 77102.6 ) | NS | 6713.36 ( 0 - 92279.94 ) | 28215.88 ( 12149.67 - 59332.34 ) | NS |

### Serum

| | GCS | | | mRS (30days) | | | LoS (30days) | | | Barthel (30 days) | | |
|---|---|---|---|---|---|---|---|---|---|---|---|---|
| Biomarker | Normal (GCS = 15) n = 15 | Abnormal (GCS < 15) n = 11 | p value | Good Outcome (mRS<3) n = 16 | Poor Outcome (mRS>2) n = 15 | p value | Good recovery (LoS > 2) n 18= | Poor Recovery (LoS =2) n = 11 | p value | Independent (score =20) n = 13 | High dependency (score <20) n = 18 | p value |
| IL-1α | 0.37 (0.21 - 0.48) | 0.43 (0.24 - 0.5) | NS | 0.31 (0.2 - 0.44) | 0.46 (0.28 - 0.53) | NS | 0.33 (0.21 - 0.48) | 0.46 (0.33 - 0.53) | NS | 0.37 (0.19 - 0.46) | 0.43 (0.27 - 0.51) | 0.36 |
| IL-1β | 0.04 (0.03 - 0.04) | 0.04 (0.03 - 0.05) | NS | 0.04 (0.03 - 0.04) | 0.04 (0.03 - 0.05) | NS | 0.04 (0.03 - 0.04) | 0.04 (0.03 - 0.05) | NS | 0.04 (0.03 - 0.04) | 0.04 (0.03 - 0.05) | 0.4 |
| IL-1RA | 6.91 (4.85 - 9.79) | 8.19 (4.92 - 10.94) | NS | 6.82 (4.7 - 7.99) | 10.2 (5.36 - 11.4) | NS | 5.76 (4.28 - 7.84) | 10.88 (6.26 - 11.4) | 0.02 | 5.75 (4.56 - 7.6) | 9.52 (5.63 - 11.14) | 0.04 |
| IL-2Rα | 1.92 (1.39 - 2.74) | 2.08 (1.1 - 2.69) | NS | 2.09 (1.38 - 2.62) | 1.9 (1.15 - 2.83) | NS | 2.05 (1.23 - 2.58) | 1.9 (1.29 - 3.01) | NS | 1.92 (1.44 - 2.98) | 2.08 (1.2 - 2.69) | 0.54 |
| IL-4 | 0.05 (0.03 - 0.07) | 0.05 (0.04 - 0.06) | NS | 0.05 (0.03 - 0.07) | 0.05 (0.05 - 0.06) | NS | 0.05 (0.03 - 0.07) | 0.05 (0.05 - 0.06) | NS | 0.04 (0.03 - 0.07) | 0.05 (0.05 - 0.06) | 0.22 |
| IL-6 | 0.07 (0.06 - 0.1) | 0.25 (0.11 - 1.18) | 0 | 0.07 (0.05 - 0.09) | 0.19 (0.09 - 0.71) | 0 | 0.07 (0.06 - 0.09) | 0.23 (0.13 - 0.71) | 0 | 0.08 (0.05 - 0.09) | 0.14 (0.07 - 0.6) | 0.05 |
| IL-8 | 0.25 (0.21 - 0.37) | 0.37 (0.26 - 0.51) | NS | 0.26 (0.21 - 0.38) | 0.26 (0.22 - 0.45) | NS | 0.25 (0.2 - 0.38) | 0.3 (0.22 - 0.47) | NS | 0.28 (0.22 - 0.39) | 0.25 (0.22 - 0.43) | 0.88 |
| IL-9 | 12.35 (11.15 - 18.38) | 11.67 (8.69 - 15.04) | NS | 11.61 (9.85 - 16.46) | 11.95 (8.73 - 15.09) | NS | 11.62 (9.81 - 15.28) | 13.12 (10.02 - 17.08) | NS | 11.67 (9.86 - 16.28) | 11.7 (8.69 - 15.45) | 0.82 |
| IL-10 | 0.12 (0.08 - 0.14) | 0.08 (0.07 - 0.12) | NS | 0.12 (0.09 - 0.14) | 0.08 (0.03 - 0.1) | NS | 0.11 (0.07 - 0.14) | 0.08 (0.05 - 0.11) | NS | 0.12 (0.07 - 0.15) | 0.08 (0.04 - 0.12) | 0.17 |
| IL-12-p40- | 1.69 (1.29 - 2.32) | 1.41 (0.98 - 2.62) | NS | 1.71 (1.25 - 2.1) | 1.76 (0.99 - 2.71) | NS | 1.57 (1.19 - 2.07) | 2.02 (1.19 - 2.71) | NS | 1.74 (1.29 - 2.09) | 1.52 (0.98 - 2.77) | 1 |
| IL-13 | 0.04 (0.01 - 0.05) | 0.04 (0.02 - 0.06) | NS | 0.04 (0.02 - 0.06) | 0.04 (0.02 - 0.05) | NS | 0.04 (0.01 - 0.06) | 0.04 (0.02 - 0.05) | NS | 0.04 (0.02 - 0.06) | 0.04 (0.02 - 0.05) | 0.4 |
| IL-16 | 0.98 (0.76 - 1.05) | 0.99 (0.81 - 1.16) | NS | 1 (0.84 - 1.21) | 0.99 (0.85 - 1.13) | NS | 0.98 (0.69 - 1.02) | 0.99 (0.85 - 1.16) | NS | 1.01 (0.92 - 1.2) | 0.99 (0.85 - 1.18) | 0.97 |
| IL-17A | 0.24 (0.19 - 0.27) | 0.25 (0.22 - 0.29) | NS | 0.24 (0.19 - 0.27) | 0.25 (0.23 - 0.29) | NS | 0.25 (0.21 - 0.28) | 0.25 (0.22 - 0.29) | NS | 0.25 (0.2 - 0.28) | 0.25 (0.2 - 0.28) | 0.76 |
| IL-18 | 1.26 (0.98 - 1.57) | 1.06 (0.85 - 1.37) | NS | 1.39 (0.99 - 1.58) | 1.08 (0.95 - 1.55) | NS | 1.22 (0.97 - 1.58) | 1.11 (0.95 - 1.55) | NS | 1.26 (0.96 - 1.54) | 1.13 (1 - 1.63) | 0.97 |
| TNF | 1.09 (0.88 - 1.72) | 1.31 (0.82 - 1.64) | NS | 1.08 (0.83 - 1.58) | 1.26 (0.83 - 1.62) | NS | 1.14 (0.86 - 1.5) | 1.29 (0.83 - 1.7) | NS | 1.08 (0.91 - 1.56) | 1.21 (0.82 - 1.64) | 0.68 |
| LTα | 27.09 (18.48 - 57.22) | 26.05 (16.5 - 40.49) | NS | 21.39 (16.15 - 37.24) | 27.53 (17.55 - 43.39) | NS | 22.01 (16.75 - 40.66) | 28.33 (17.06 - 43.7) | NS | 23.51 (16.29 - 28.1) | 27.09 (16.27 - 43.88) | 0.88 |
| IFN-α2 | 0.1 (0.08 - 0.13) | 0.11 (0.07 - 0.15) | NS | 0.11 (0.08 - 0.15) | 0.1 (0.07 - 0.15) | NS | 0.11 (0.08 - 0.14) | 0.1 (0.07 - 0.14) | NS | 0.11 (0.09 - 0.14) | 0.09 (0.07 - 0.16) | 0.71 |
| IFNγ | 0.23 (0.14 - 0.56) | 0.31 (0.27 - 0.51) | NS | 0.29 (0.17 - 0.56) | 0.28 (0.19 - 0.51) | NS | 0.29 (0.16 - 0.54) | 0.28 (0.19 - 0.52) | NS | 0.27 (0.15 - 0.55) | 0.31 (0.2 - 0.51) | 0.82 |
| CXCL1 | 17.86 (4.68 - 39.04) | 20.83 (8.11 - 26.1) | NS | 8.1 (4.41 - 28.73) | 20.28 (9.22 - 28.17) | NS | 15.04 (4.67 - 39.11) | 20.16 (5.55 - 26.62) | NS | 6.96 (3.68 - 23.95) | 19.74 (7.56 - 29.2) | 0.29 |
| CXCL9 | 4.55 (2.82 - 5.97) | 4.87 (3.92 - 7.78) | NS | 4.28 (2.9 - 6.03) | 5.02 (3.94 - 6.8) | NS | 5.32 (3.34 - 6.83) | 4.88 (3.42 - 5.61) | NS | 4.55 (3.01 - 6) | 4.89 (2.97 - 7.08) | 0.79 |
| CXCL10 | 18.75 (12.42 - 26.36) | 16.2 (14.33 - 28.85) | NS | 18.82 (14.32 - 26.26) | 15.19 (11.24 - 23.96) | NS | 19.4 (14.79 - 30.89) | 14 (10.36 - 20.57) | 0.05 | 18.9 (14.11 - 26.15) | 16 (12.49 - 24.55) | 0.62 |
| CXCL12 | 38.82 (34.44 - 56.17) | 40.33 (35.23 - 51.35) | NS | 38.93 (35.35 - 54.99) | 40.91 (33.87 - 51.12) | NS | 40.26 (34.51 - 53.82) | 38.28 (33.87 - 48.15) | NS | 38.14 (34.1 - 54.39) | 41.49 (34.51 - 51.35) | 0.97 |
| CCL2 | 0.79 (0.35 - 1.22) | 1.11 (0.96 - 1.89) | 0.05 | 1 (0.63 - 1.28) | 1 (0.8 - 1.34) | NS | 0.93 (0.63 - 1.33) | 1 (0.86 - 1.2) | NS | 0.95 (0.63 - 1.14) | 1.03 (0.79 - 1.31) | 0.47 |
| CCL3 | 0.09 (0.07 - 0.13) | 0.06 (0.05 - 0.09) | 0.03 | 0.08 (0.06 - 0.1) | 0.09 (0.05 - 0.11) | NS | 0.08 (0.07 - 0.11) | 0.09 (0.05 - 0.12) | NS | 0.08 (0.07 - 0.12) | 0.09 (0.05 - 0.11) | 0.52 |
| CCL4 | 16.88 (11.62 - 20.65) | 13.15 (10.63 - 14.77) | NS | 15.57 (11.84 - 20.4) | 12.09 (10.72 - 14.73) | NS | 14.65 (11 - 19.69) | 12.91 (11.27 - 15.56) | NS | 14.63 (11.2 - 20.14) | 12.67 (10.84 - 16.25) | 0.45 |
| CCL5 | 152.05 (119.57 - 224.1) | 197.56 (163.5 - 246.69) | NS | 185.46 (135.47 - 217.17) | 176.17 (126.29 - 241.21) | NS | 182.04 (142.15 - 209.13) | 153 (118.82 - 241.21) | NS | 177.41 (139.96 - 210.23) | 186.67 (125.01 - 243.37) | 0.82 |
| CCL27 | 32.06 (27.99 - 36.99) | 29.04 (16.97 - 38.09) | NS | 28.52 (26 - 36.83) | 32.47 (19.74 - 36) | NS | 33.62 (28.34 - 39.53) | 27.6 (18.64 - 34.22) | NS | 28.35 (27.63 - 34.37) | 32.88 (19.54 - 37.17) | 0.65 |
| LIF | 0.78 (0.41 - 0.87) | 0.88 (0.68 - 1.06) | NS | 0.71 (0.4 - 0.91) | 0.86 (0.74 - 1.02) | NS | 0.71 (0.41 - 0.95) | 0.86 (0.8 - 1.02) | NS | 0.78 (0.41 - 0.89) | 0.84 (0.57 - 1.04) | 0.4 |
| M-CSF | 0.52 (0.38 - 0.58) | 0.41 (0.29 - 0.84) | NS | 0.4 (0.34 - 0.54) | 0.55 (0.33 - 0.92) | NS | 0.4 (0.3 - 0.56) | 0.75 (0.48 - 0.97) | 0.02 | 0.38 (0.35 - 0.53) | 0.52 (0.32 - 0.85) | 0.4 |
| MIF | 8.7 (6.27 - 11.62) | 10.34 (8.16 - 12.82) | NS | 9.77 (6.69 - 12.41) | 9.33 (6.87 - 16.99) | NS | 8.14 (6.26 - 11.99) | 9.94 (7.5 - 16.99) | NS | 9.53 (6.29 - 12.36) | 9.53 (7.38 - 14.76) | 0.6 |
| PDGF-bb | 44.31 (26.02 - 52.06) | 35.49 (28.06 - 76.94) | NS | 38.95 (23.86 - 56.24) | 44.09 (28.14 - 52.89) | NS | 38.58 (28.81 - 52.76) | 49.21 (28.06 - 66.3) | NS | 44.31 (23.94 - 56.12) | 41.66 (28.13 - 55.99) | 0.97 |
| SCF | 2.23 (1.93 - 2.64) | 1.96 (1.65 - 2.42) | NS | 2.21 (1.91 - 2.63) | 2.19 (1.77 - 2.5) | NS | 2.21 (1.86 - 2.65) | 2.24 (1.9 - 2.5) | NS | 2.22 (1.76 - 2.65) | 2.15 (1.87 - 2.53) | 0.82 |
| SCGF-β | 1167.47 (986.29 - 1440.66) | 950.1 (804.75 - 1210.19) | NS | 1087.63 (929.32 - 1380.94) | 1152.81 (919.43 - 1271.19) | NS | 1110.84 (866.74 - 1307.11) | 1189.72 (1001.74 - 1378.51) | NS | 1062.7 (852.64 - 1321.23) | 1155.29 (984.53 - 1277.29) | 0.52 |
| TRAIL | 0.67 (0.17 - 1.58) | 0.23 (0.14 - 1.01) | NS | 1.15 (0.31 - 1.56) | 0.22 (0.1 - 0.92) | 0.02 | 0.91 (0.3 - 1.49) | 0.17 (0.1 - 0.92) | 0.03 | 1.15 (0.2 - 1.54) | 0.28 (0.11 - 1.17) | 0.1 |
| Eotaxin | 2.55 (1.92 - 3.13) | 2.18 (1.25 - 2.5) | NS | 2.75 (2.14 - 3.69) | 1.86 (1.3 - 2.4) | 0.03 | 2.63 (2.18 - 3.53) | 1.62 (1.06 - 2.29) | 0 | 2.61 (2.04 - 3.18) | 2.18 (1.41 - 2.6) | 0.3 |
| FGF-basic | 0.9 (0.51 - 1.32) | 0.94 (0.45 - 1.09) | NS | 0.71 (0.5 - 1.15) | 0.97 (0.48 - 1.1) | NS | 0.9 (0.51 - 1.32) | 0.94 (0.47 - 1.07) | NS | 0.72 (0.51 - 1.07) | 0.95 (0.43 - 1.1) | 0.97 |
| G-CSF | 1.5 (1.08 - 4.55) | 2.19 (1.77 - 2.67) | NS | 1.33 (1.03 - 2.78) | 2.34 (1.8 - 4.04) | NS | 1.51 (1.06 - 4.07) | 2.34 (1.87 - 3.81) | NS | 1.2 (1.05 - 2.83) | 2.16 (1.61 - 3.84) | 0.45 |
| HGF | 15.09 (10.66 - 24.51) | 13.2 (10.91 - 19.84) | NS | 16.85 (12.94 - 24.34) | 12.88 (9.87 - 18.9) | NS | 14.24 (11.67 - 24.16) | 13.61 (9.87 - 22.66) | NS | 15.09 (11.6 - 24.16) | 13.2 (10.91 - 22.93) | 0.79 |
| GFAP-pg-mL | 970.99 (384.07 - 2350.47) | 6651.16 (3873.34 - 10367.41) | 0.01 | 917.88 (263.57 - 2227.68) | 5517.62 (3479.58 - 11547.01) | 0 | 951.6 (321.39 - 3577.79) | 6628.15 (2282.5 - 12417.61) | 0.01 | 903.55 (198.53 - 1726.09) | 3878.21 (2354.64 - 10367.41) | 0 |
| NFL-pg-mL | 33.15 (25.64 - 151.57) | 161.18 (90.64 - 338.39) | 0.03 | 32.9 (21.55 - 86.83) | 173.15 (99.93 - 323.86) | 0 | 33.25 (23.9 - 99.93) | 192.63 (155.64 - 375.7) | 0 | 33.15 (19.71 - 84.91) | 161.18 (81.3 - 260.64) | 0 |
| Tau-pg-mL | 2.29 (1.46 - 3.91) | 10.66 (6.98 - 17.16) | 0 | 1.89 (1.41 - 3.13) | 9.18 (5.77 - 14.87) | 0 | 2.14 (1.45 - 4.87) | 7.88 (5.77 - 13.61) | 0 | 1.98 (1.48 - 2.81) | 8.61 (5.59 - 13.78) | 0 |
| UCH-L1-pg-mL | 28.22 (22.76 - 53.5) | 178.07 (135.09 - 241.51) | 0 | 32.19 (25.29 - 70.09) | 154.82 (85.83 - 231.57) | 0.01 | 40.43 (26.95 - 106.63) | 154.82 (79.97 - 263.91) | 0.05 | 33.53 (20.76 - 101.24) | 113.4 (58.15 - 200.09) | 0.04 |

Summary statistics are median (IQR). NS = *P value* ≥0.05, acute period = 0-9 days post-randomisation

**Supplementary Table 2:** KEGG enrichment pathway analysis of the 9 mediators associated with clinical severity

| Category | Term | Count | P-Value | Benjamini | Fold Enrichment | Bonferroni | log10_benjamini |
|---|---|---|---|---|---|---|---|
| KEGG_PATHWAY | Cytokine-cytokine receptor interaction | 8 | 4.59E-10 | 3.26E-08 | 25.46 | 3.26E-08 | 7.4867824 |
| KEGG_PATHWAY | Malaria | 4 | 0.0000104 | 0.000369 | 75.86 | 0.000737 | 3.432973634 |
| KEGG_PATHWAY | Rheumatoid arthritis | 4 | 0.0000719 | 0.0013 | 39.93 | 0.00509 | 2.886056648 |
| KEGG_PATHWAY | Viral protein interaction with cytokine and cytokine receptor | 4 | 0.0000838 | 0.0013 | 37.93 | 0.00593 | 2.886056648 |
| KEGG_PATHWAY | Chagas disease | 4 | 0.0000915 | 0.0013 | 36.82 | 0.00648 | 2.886056648 |
| KEGG_PATHWAY | JAK-STAT signaling pathway | 4 | 0.00039 | 0.00431 | 22.58 | 0.0273 | 2.36552273 |
| KEGG_PATHWAY | Influenza A | 4 | 0.000425 | 0.00431 | 21.92 | 0.0298 | 2.36552273 |
| KEGG_PATHWAY | African trypanosomiasis | 3 | 0.000504 | 0.00447 | 76.88 | 0.0351 | 2.349692477 |
| KEGG_PATHWAY | Lipid and atherosclerosis | 4 | 0.000815 | 0.00643 | 17.56 | 0.0563 | 2.191789027 |
| KEGG_PATHWAY | Inflammatory bowel disease | 3 | 0.0016 | 0.0114 | 43.10 | 0.108 | 1.943095149 |
| KEGG_PATHWAY | IL-17 signaling pathway | 3 | 0.00329 | 0.0212 | 29.94 | 0.208 | 1.673664139 |
| KEGG_PATHWAY | Toll-like receptor signaling pathway | 3 | 0.0043 | 0.0255 | 26.10 | 0.264 | 1.59345982 |
| KEGG_PATHWAY | Herpes simplex virus 1 infection | 3 | 0.0116 | 0.059 | 15.63 | 0.564 | 1.229147988 |
| KEGG_PATHWAY | Tuberculosis | 3 | 0.0116 | 0.059 | 15.63 | 0.564 | 1.229147988 |
| KEGG_PATHWAY | Chemokine signaling pathway | 3 | 0.013 | 0.0616 | 14.74 | 0.606 | 1.210419288 |
